# Rfam 15: RNA families database in 2025

**DOI:** 10.1101/2024.09.23.614430

**Authors:** Nancy Ontiveros, Emma Cooke, Eric P. Nawrocki, Sandra Triebel, Manja Marz, Elena Rivas, Sam Griffiths-Jones, Anton I. Petrov, Alex Bateman, Blake Sweeney

**Affiliations:** European Molecular Biology Laboratory, Wellcome Genome Campus, European Bioinformatics Institute, Hinxton, Cambridge, CB10 1SD, UK; SciBite, Cambridge, UK; National Center for Biotechnology Information, U.S. National Library of Medicine, National Institutes of Health, Bethesda, MD, 20894, USA (EPN); RNA Bioinformatics and High-Throughput Analysis, Friedrich Schiller University Jena, 07743, Jena, Germany; European Virus Bioinformatics Center, Friedrich Schiller University Jena, 07743, Jena, Germany; Department of Molecular and Cellular Biology, Harvard University, Cambridge, 02138, MA, USA; School of Biological Sciences, Faculty of Medicine, Biology and Health, Michael Smith Building, The University of Manchester, Manchester M13 9GB, UK; Riboscope Ltd., Cambridge, CB1 1AH, UK

**Keywords:** Non-coding RNA, Viruses, microRNA, RNA 3D Structure, Bioinformatics, Database

## Abstract

The Rfam database, a widely-used repository of non-coding RNA (ncRNA) families, has undergone significant updates in release 15.0. This paper introduces major improvements, including the expansion of Rfamseq to 26, 106 genomes, a 76% increase, incorporating the latest UniProt reference proteomes and additional viral genomes. Sixty-five RNA families were enhanced using experimentally determined 3D structures, improving the accuracy of consensus secondary structures and annotations. R-scape covariation analysis was used to refine structural predictions in 26 families. Gene Ontology and Sequence Ontology annotations were comprehensively updated, increasing GO term coverage to 75% of families. The release adds 14 new Hepatitis C Virus RNA families and completes microRNA family synchronisation with miRBase, resulting in 1, 603 microRNA families. New data types, including FULL alignments, have been implemented. Integration with APICURON for improved curator attribution and multiple website enhancements further improve user experience. These updates significantly expand Rfam’s coverage and improve annotation quality, reinforcing its critical role in RNA research, genome annotation, and the development of machine learning models. Rfam is freely available at https://rfam.org.

**Graphical Abstract:** Rfam has undergone a major update with the release of 15.0. We have increased the number of genomes in our sequence database Rfamseq by 75%, completed the synchronisation with miRBase and improved 65 families using 3D structures.

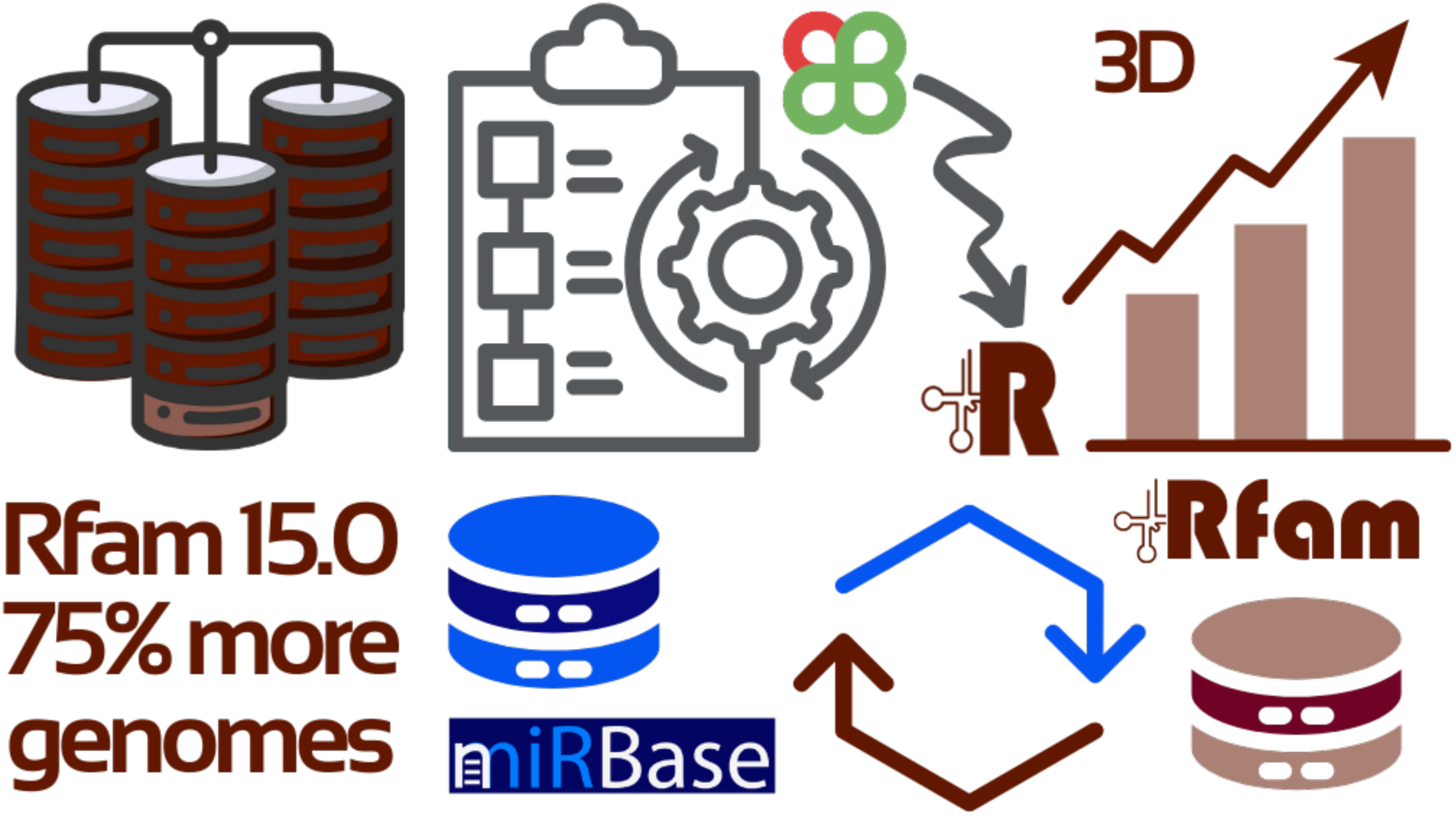

## Introduction

The Rfam database was established in 2002 (1) in order to provide a central repository of non-coding RNA (ncRNA) families for genomic annotations. Each family is represented by three key components: (i) a multiple sequence alignment called the SEED alignment that contains aligned sequences of homologous RNA sequences that share a consensus secondary structure, (ii) a covariance model (CM) built using the Infernal software (2) that was trained on the SEED alignment, and (ii) a set of matches that were found using the covariance model, called the set of FULL hits. The families are built manually by Rfam curators using scientific literature or alignments submitted by the community.

For each model the curators search for homologs in a sequence database, Rfamseq, and select a bit score threshold, called the gathering threshold, that separates homologous sequences from unrelated or more distantly related sequences, taking into account phylogenetic distribution of hits. The cutoff can be used by Infernal to report only sequences that score higher than this threshold when annotating genomes or searching in sequence databases.

Each major release of the Rfam database, such as 14.0 or 15.0, corresponds to a new version of Rfamseq. Prior to Rfam 13.0, Rfamseq was based on a set of sequences from the ENA database (3). However, due to the explosive growth in ENA, in Rfam 13.0 Rfamseq transitioned to a representative and reduced redundancy set of genomes produced by the UniProt team (4) for the Reference Proteomes dataset (5). The representative proteomes from UniProt are mapped to their corresponding genomes and the genomic sequence is analysed by Rfam (5). This provides users with a representative collection of genomes which have comprehensive ncRNA and protein annotations.

Rfam has been widely adopted by various scientific communities, with uses ranging from the annotation of small non-coding RNAs (ncRNAs) in genomic resources like Ensembl (6), to serving as a dataset for training machine learning models such as AlphaFold 3 (7). Additionally, Rfam is frequently used as a key reference for known non-coding RNAs. To ensure it remains current and valuable to the scientific community, we continuously review, update, and enhance the Rfam families.

In release 15.0 we have updated and expanded Rfamseq, improved families using 3D structures and R-scape (8), improved Gene Ontology (9) and Sequence Ontology (10) annotations and completed the synchronisation of miRBase (11) microRNAs into Rfam.

## New Rfamseq database

Rfam 15 is based on the 2024_03 release of UniProt reference proteomes that includes 23, 158 genomes (see Figure 1 showing the taxonomic distribution). We attempted to download all components of the genomes used to build the proteome set. In some cases, the specified genome was replaced by a newer version, which was used instead. In a very few cases, less than 50, all versions of the genome were either outdated or removed. In those cases we did not fetch any genome.

**Figure 1.**
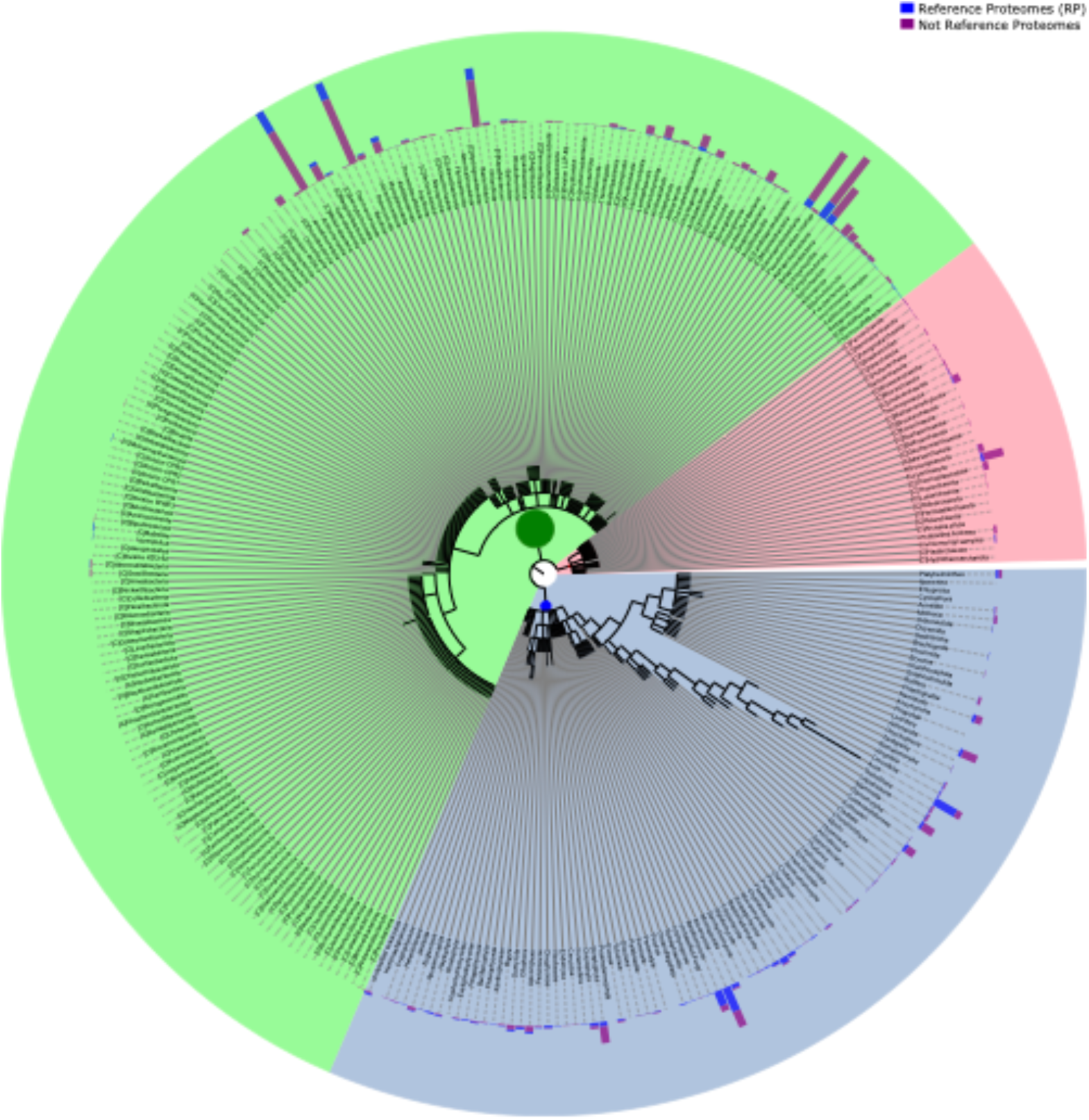
The taxonomic distribution of cellular organisms in Rfamseq 15 organised by the phylogenetic kingdom. Shown on the outside is the number of reference and non-reference proteomes in each lineage. Each coloured section indicates the kingdom with bacteria (green), archaea (pink) and eukaryotes (blue).

In addition to the genomes in the UniProt set, we also fetched all viral genomes from the protein information resource (12) at 75% percent identity for an additional 2, 985 viral genomes. This set was added because during our viral work, described below, we found that the UniProt reference proteome work under reports the diversity of viral genomes. This has led to some viral Rfam families having very few or no FULL hits. Rfam would prefer that all families have at least one match outside of the SEED sequences. While using an additional set of viral genomes leads to a large increase in the number of genomes, the total size of Rfamseq is dominated by cellular organisms as <1% of the nucleotides come from a viral genome. Table 1 shows the comparison of Rfamseq in the last major release Rfam 14.0 vs Rfam 15.0.

**Table 1.**
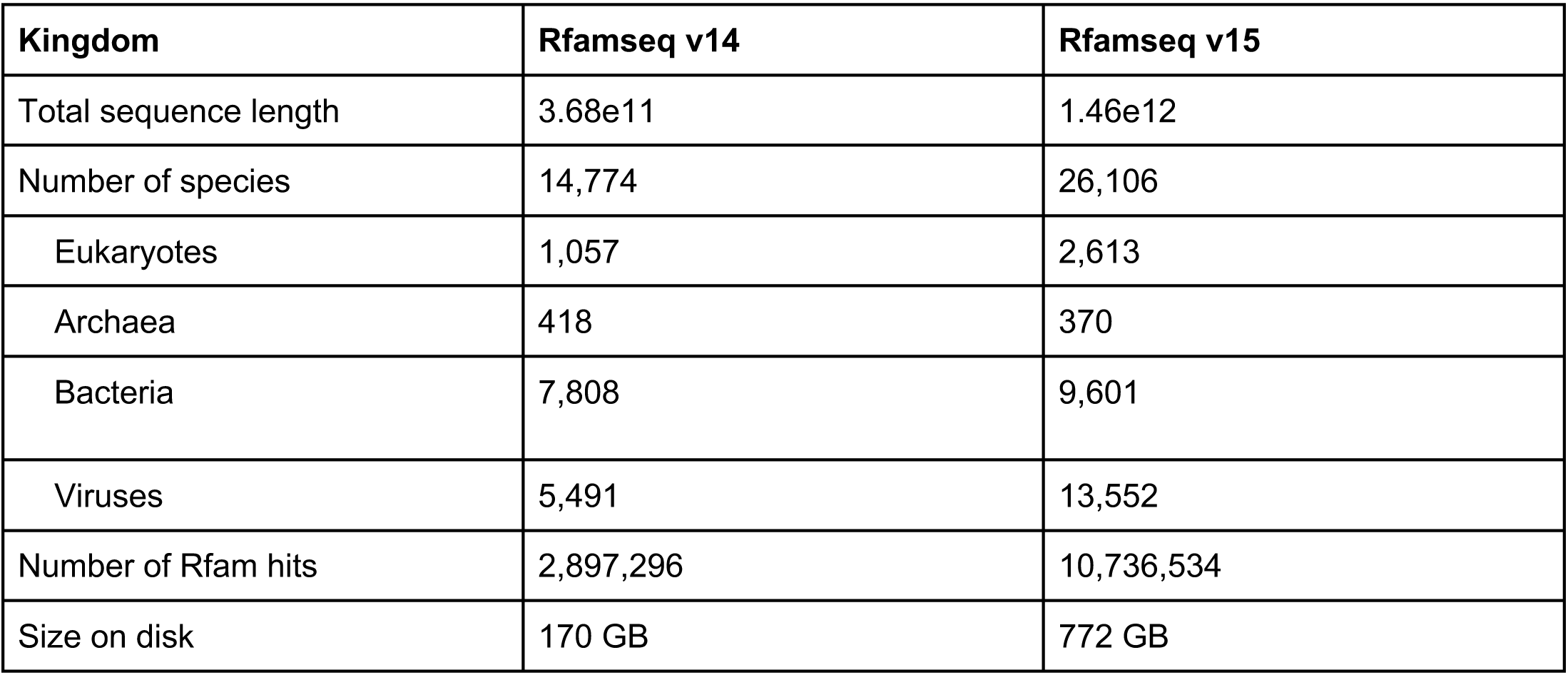
The number of genomes in Rfamseq corresponding to release 14.0 and 15.0. The size on disk is the uncompressed size of all sequences in Rfamseq.

| Kingdom | Rfamseq v14 | Rfamseq v15 |
| --- | --- | --- |
| Total sequence length | 3.68e11 | 1.46e12 |
| Number of species | 14,774 | 26,106 |
| Eukaryotes | 1,057 | 2,613 |
| Archaea | 418 | 370 |
| Bacteria | 7,808 | 9,601 |
| Viruses | 5,491 | 13,552 |
| Number of Rfam hits | 2,897,296 | 10,736,534 |
| Size on disk | 170 GB | 772 GB |

Once Rfamseq was updated all families were searched against the updated database and the matches were collected. Many older or larger families have not been rescanned in years due to technical pipeline limitations, for example, the 5S rRNA (RF00001) has not had its matches updated since 2014.

Once all families were updated, we compared the matches of families in 14.3, which was the last published version, and 15.0 to determine how Rfam has changed. Of the 3, 431 families in common between 14.3 and 15.0, 98 had no FULL hits in 14.3 and when moving to 15.0, 23 of these families gained at least one hit while 21 lost all hits, leading to 96 families in 15.0 without FULL hits. Families without FULL hits occur because either the curator-selected gathering threshold of the family is too high, or if Rfamseq does not contain any similar sequences. Future work will look to see if these families require updating the threshold or if additional genomes need to be added to Rfamseq.

Of the 3, 302 families with hits in both releases, the average family grew by 166%, with 2, 335 gaining at least one hit, 547 losing hits and 420 with no change in the number of hits. Examining the families with an increase shows that there are 26 families with a greater than 10x growth. These families are primarily microRNA families matching to plant genomes. While, as described below, these families have been synchronised with miRBase, we are revisiting these families to see if they need additional future updates in the light of these additional hits, or if the genomes are poorly assembled and should be excluded from Rfamseq.

Overall the update to Rfamseq has led to a considerable change in all Rfam families. This will provide the community with a consistent and up-to-date dataset to reuse. Future updates to Rfamseq will seek to maximise the coverage of known organisms.

## Improving existing Rfam families

### Using RNA 3D structures to revise Rfam secondary structures

Rfam families are based on multiple sequence alignments annotated with consensus secondary structures that indicate base pairing. The accuracy and completeness of the consensus secondary structure is of critical importance as it guides the SEED alignment, informs the Infernal CM, and is used for training and benchmarking of software for RNA 2D and 3D structure prediction, including AlphaFold 3 (7) and R-scape (8).

Most of the Rfam families with known 3D structure were created prior to the 3D structure determination using less accurate, predicted secondary structures. For example, while the FMN riboswitch family (RF00050) correctly captured 5 helices of the FMN riboswitch, it did not include 1 helix, 2 pseudoknots, and several key base pairings (Figure 2B). As such, one major goal of this project was to include the pseudoknots which are observed in 3D structures into the Rfam families.

**Figure 2.**
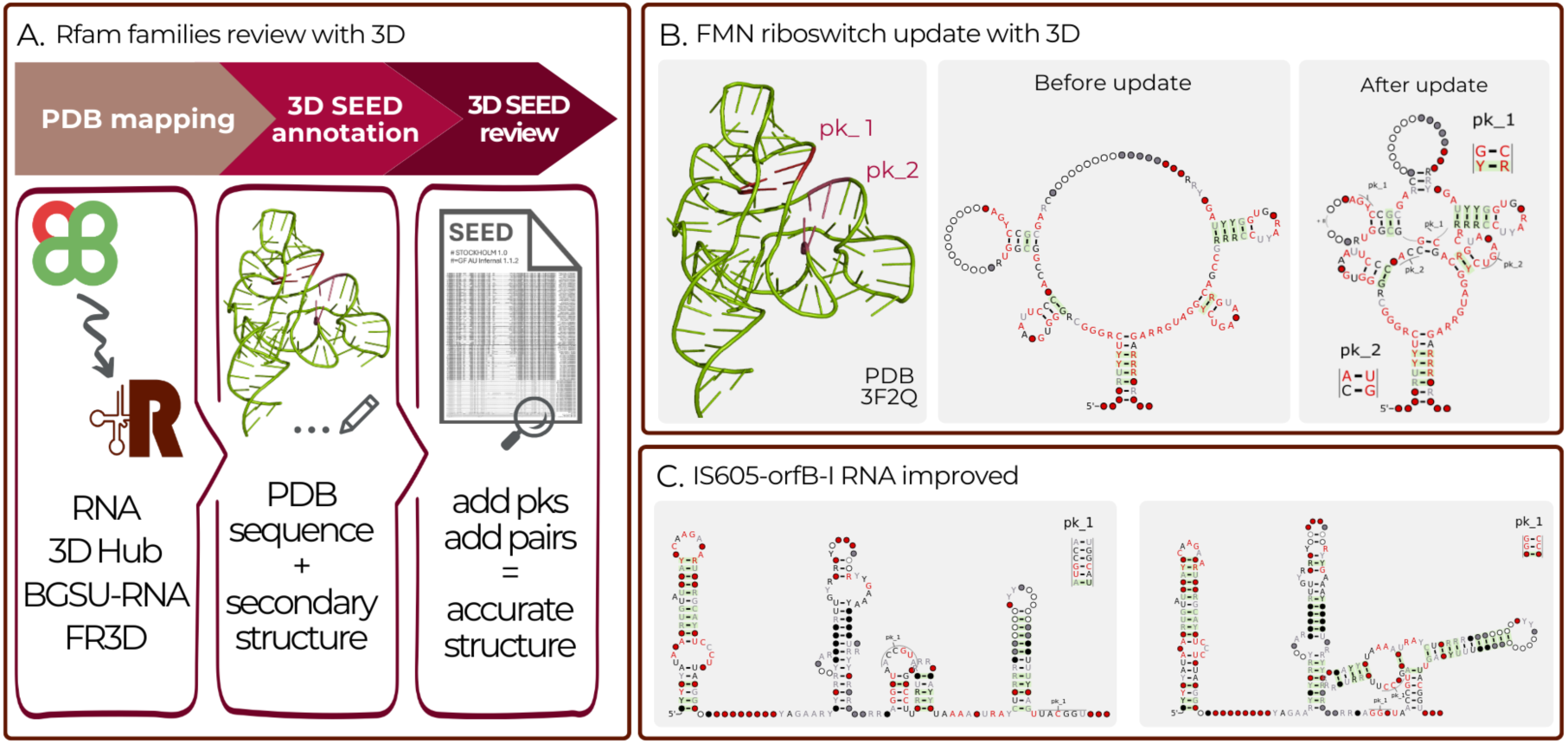
A) Pipeline for reviewing Rfam families using experimentally determined RNA 3D structures from PDB. B) Example of the FMN riboswitch (RF00050) before and after improvements with a 3D structure. Improvements included adding 2 pseudoknots and creating a missing helicees. C) A summary of how the IS605-orfB-I RNA (RF03065) family was reviewed and improved using R-scape.

To aid the Rfam curators with updating Rfam alignments an automated pipeline was developed to map RNA sequences and secondary structures of experimentally determined RNA 3D structures to Rfam SEED alignments. Every week Rfam families are mapped to the latest set of RNA chains from PDB (13) using the Infernal *cmscan* program. For each Rfam family with one or more matching structures, the sequences and secondary structures are iteratively added to the alignment in the Stockholm format using Infernal’s *cmalign* program. The base pairing information, as determined by FR3D (14), is also included in the alignment as an additional GR annotation line (one per 3D structure). The pipeline is automatically executed weekly and the Rfam curators receive notifications of newly mapped structures (Figure 2A).

The resulting alignments undergo manual review to determine whether the family consensus secondary structure is consistent with the base pair information from PDB structures. The structure and annotated based pairs are always kept in the alignment, even in cases where the structure is not used to update the consensus secondary structure, e.g. low resolution structure or non-canonical base pairs. Additional annotations are also included in the Stockholm files, such as RNA structural elements. For example P1, P2, and P3 domains are included in the Pistol ribozyme RF02679, while the SAM riboswitch (RF00162) now includes a kink-turn annotation.

Our pipeline detects 143 Rfam families which match at least one 3D structure. We began updating families with 3D structures in release 14.5 and since then Rfam has updated 65 families, with 298 chains from 3D structures. This includes well known families such as the SAM riboswitch (RF00162), the FMN riboswitch (RF00050) and microRNA 16 precursor (RF00254). The updates included the addition of pseudoknots (PK) in 42 of the 65 families, of which 6 had 2 PK, and 1 had 3. These features include ligand binding sites for riboswitches (28 annotations), locations of well known motifs such kink-turns (8 annotations), and structural regions such as helices and domains (60 annotations). A detailed table showing which families were updated with which 3D structures and can also be found online at https://rfam.org/3d. Rfam curators work to keep families as up-to-date with the latest structures as possible, which has led to 11 families being updated more than once since release 14.5.

The review of structures is part of the ongoing curation process, and additional families will be improved in future releases. The pipeline is implemented in Python and is available at https://github.com/Rfam/rfam-3d-seed-alignments and the weekly updates are found in the file pdb_full_region.txt.gz located in the preview section of the FTP archive (http://ftp.ebi.ac.uk/pub/databases/Rfam/.preview/).

### Using R-scape to improve Rfam consensus secondary structures

Since version 13.0 (5) the Rfam website included the results of the R-scape analysis (8) to help users evaluate consensus secondary structures for each Rfam family, as well as alternative secondary structures generated using the R-scape CaCoFold algorithm. CaCoFold evaluates all possible consensus secondary structures that are consistent with multiple sequence alignment and can propose a structure maximising the number of base pairs with statistically significant covariation (15).

In release 14.9, the R-scape software was used to systematically review all Rfam families and identify those that could be enhanced using R-scape CaCoFold structures. We examined all families with an increased number of covarying base pairs and selected 26 families for updates. These families are listed in Table 2. Shown in Figure 2C is an example of a family, IS605-orfB-I, before and after improvement.

**Table 2:**
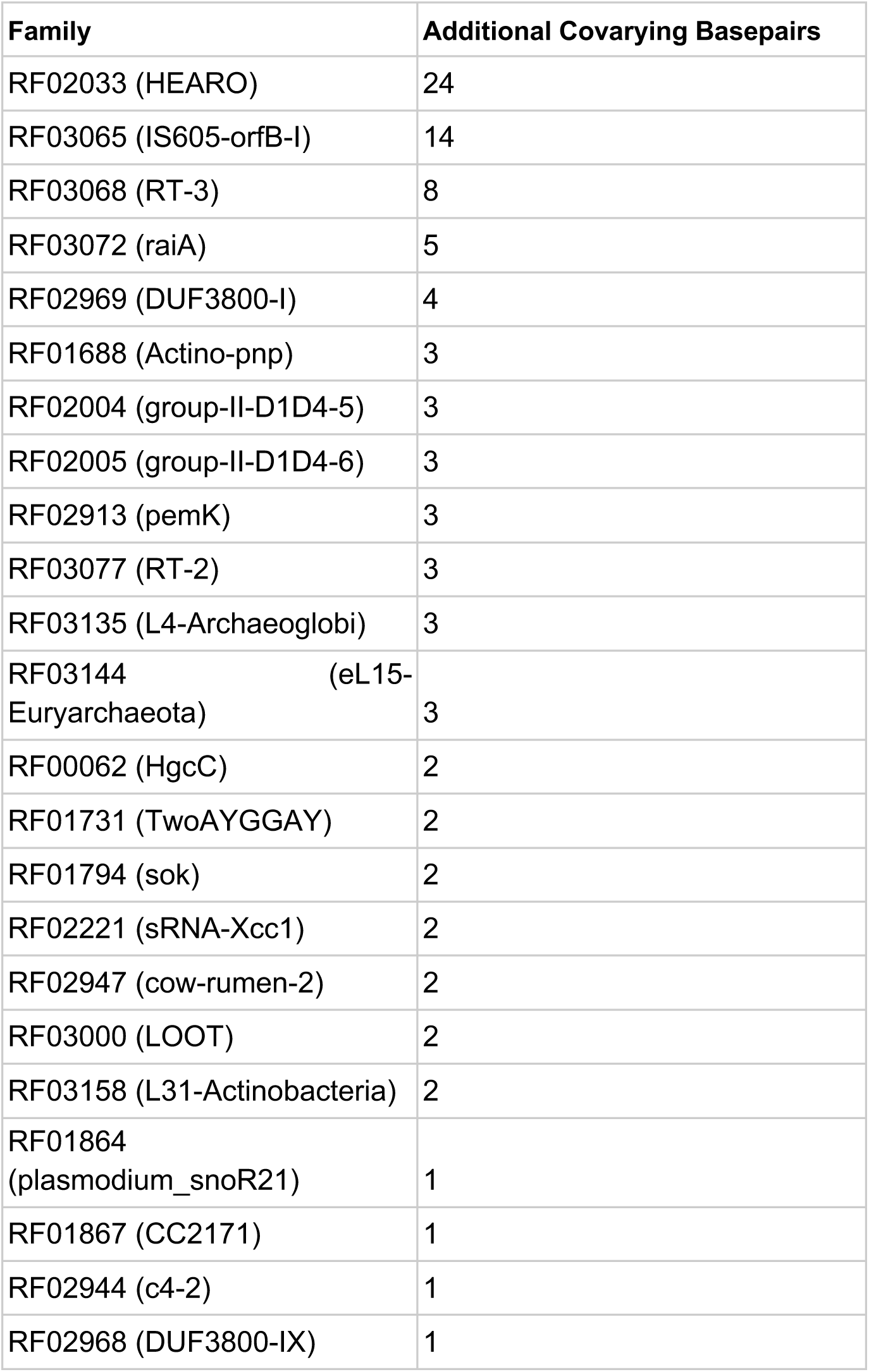
Summary of the R-scape based improvements. This shows the families which were improved by using R-scape CaCoFold structures.

## Gene and Sequence Ontology annotations

Rfam is commonly used as a source of functional information for non-coding RNAs. This takes several forms, from users reading our curated descriptions and Wikipedia articles to understand the role of an ncRNA, to using the Gene Ontology (GO) and Sequence Ontology (SO) annotations that Rfam curators provide to understand the role of an ncRNA. Rfam is the largest source of GO annotations with over 10 million sequences having an Rfam based GO annotation. In addition to providing information to human users, Rfam is also used in training Large Language Models (LLM). LLMs such as ChatGPT and Claude have clearly been trained on Rfam data and are able to output entries in the formats, Stockholm and DESC, that Rfam uses. Thus providing a completely new way for scientists to access the information found in Rfam.

The Gene Ontology is a resource to classify and provide functional information for biomolecules using structured annotations and ontologies (9). Similarly the Sequence Ontology provides an ontology and structured annotations for the types of biomolecules (10). Both resources are continuously updated to better reflect scientific knowledge. For example, since the last publication of Rfam the SO gained several terms specific to the location of rRNA, e.g. cytosolic_rRNA (SO:0002343) and obsoleted the more generic rRNA terms.

Rfam provides GO and SO annotations for families, however, these annotations are not regularly reviewed to stay up-to-date with the latest changes in the ontologies. As an example, prior to 15.0 the rRNA families used obsoleted SO terms. For 15.0 we manually reviewed all annotations and updated them to better reflect changes in the GO and SO, as well as improved the specificity of annotations where possible.

We ensured that all families had at least one up-to-date SO term and the annotation was as specific as possible. For GO terms, we ensured the terms were up-to-date and added related terms where possible. A few examples of the changes we made were to 1) ensure all snoRNAs had a snoRNA specific SO term and included an RNA processing GO term; 2) add the sodium ion binding term to the NA+ riboswitch; and 3) updated families with the generic mature_transcript SO term to the more specific ncRNA SO term. Of the 3, 431 common families between 14.3 and 15.0 2, 084 had a GO term in 14.3 while 2, 426 did in 15.0. For all families in 15.0 we now have at least one GO term for 3, 157, which covers 75% of all families, an increase from 60% in 14.3. These changes increased the total number of GO terms from 3, 752 to 4, 446. As these updates propagate through the community, we expect these changes to lead to millions more sequences annotated with functional information.

## Viral RNAs

Starting with Rfam 14.3 we collaborated with the Marz group to develop a new workflow for viral RNA families (16). This workflow is based upon whole genome alignments of viruses and led to the creation of *Flaviviridae* and *Coronaviridae* families as described previously (16). Since the last publication we have continued to use this workflow and have added *Hepatitis C Virus* (HCV) (17) families. We included 14 new families and removed 4 outdated families. A schematic of the new families can be seen in Figure 3. These families include structures found in both the non-coding and coding regions of the viral alignments. All viral families organised by viral clade can be seen at https://rfam.org/viruses. We plan to continue to use the viral pipeline and improve the representation of viral families in Rfam and expect to focus on HDV and pestivirus families in the near future.

**Figure 3:**
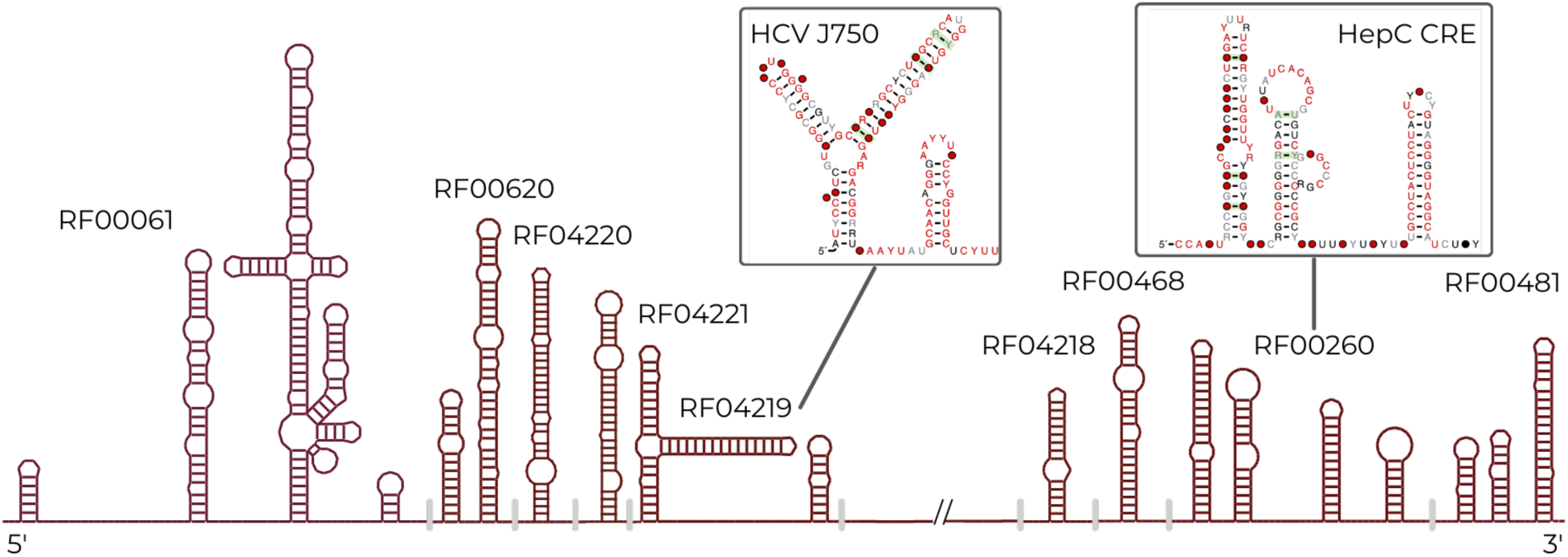
A schematic of the HCV viral families Rfam now contains. These families were built from whole genome alignments, as described in (17). The figure shows their relative locations in the HCV genomic alignment and shown in the inset are two of the new HCV families.

## MicroRNAs

MicroRNAs are an important class of ncRNA that regulate gene expression in animals and plants, with many microRNAs being implicated in disease. For example, the mir-17∼92 cluster (18) and mir-155 (19) are amongst a number of microRNAs which act as oncogenes. Understanding the evolutionary and family relationships of microRNAs across species allows the transfer of annotation, for example from model organisms to humans and vice versa.

Since early releases, Rfam has included microRNA families, e.g let-7 (20) was deposited in 2002 for Rfam 1.1. However, they were not subject to regular review and were not coordinated with miRBase (11), an authoritative resource for microRNA annotation that assigns identifiers for microRNA genes and sequences. As of Rfam 13.0, out of 1, 983 microRNA families found in miRBase v21, only 28% matched one or more of the 529 Rfam microRNA families.

Starting in Rfam 14, we began systematically reviewing microRNA families with the goal to synchronise Rfam microRNA families with miRBase. To this end, we worked to ensure that the SEED alignments of all microRNA families in Rfam were built from microRNA sequences that are tracked in the miRBase database. Multiple sequence alignments of microRNA sequences were extracted from miRBase and mapped to Rfam accessions. Each sequence was assigned a RNAcentral Unique RNA Sequence (URS) (21) identifier to remove sequence redundancy and represent only distinct sequences for each species. For each alignment a covariance model was built using Infernal and used to search the Rfamseq database. Bit score thresholds for each model (known as gathering thresholds (16)) were manually curated. These thresholds enable automatic and accurate genome annotation using Rfam microRNA families with Infernal.

Pre-existing Rfam microRNA families were reviewed and replaced with miRBase-derived SEED alignments where possible. New families were created when Rfam did not already represent the microRNA sequences. MicroRNA families found in Rfam that did not match miRBase were reviewed and updated or deleted. In cases where the miRBase alignment matched several Rfam families we manually inspected the alignment and families and split and merged families and built clans of families as appropriate.

Rfam release 15.0 marks the completion of the synchronisation process, as we have processed all microRNA alignments from miRBase. For roughly 200 miRBase-derived alignments, we determined that the family was not suitable for inclusion into Rfam. In most cases this was because there was a single unique sequence in the miRBase alignment. A small fraction of the alignments which have not been added may be removed from miRBase in the future. Table 3 shows the summary of changes since release 14.3, and a list of all microRNA families is found in Supplementary information in Table S1. All miRBase microRNA sequences that are not represented in updated Rfam families will be periodically and systematically reviewed in a cycle of improvements and curation of both Rfam and miRBase. Rfam and miRBase will continue to synchronise our resources as new microRNAs and new microRNA alignments are available, with miRBase acting as the repository of microRNA sequences, and Rfam as the authority on the grouping of those sequences into families. Rfam microRNA family classifications will be available through both resources.

**Table 3:**
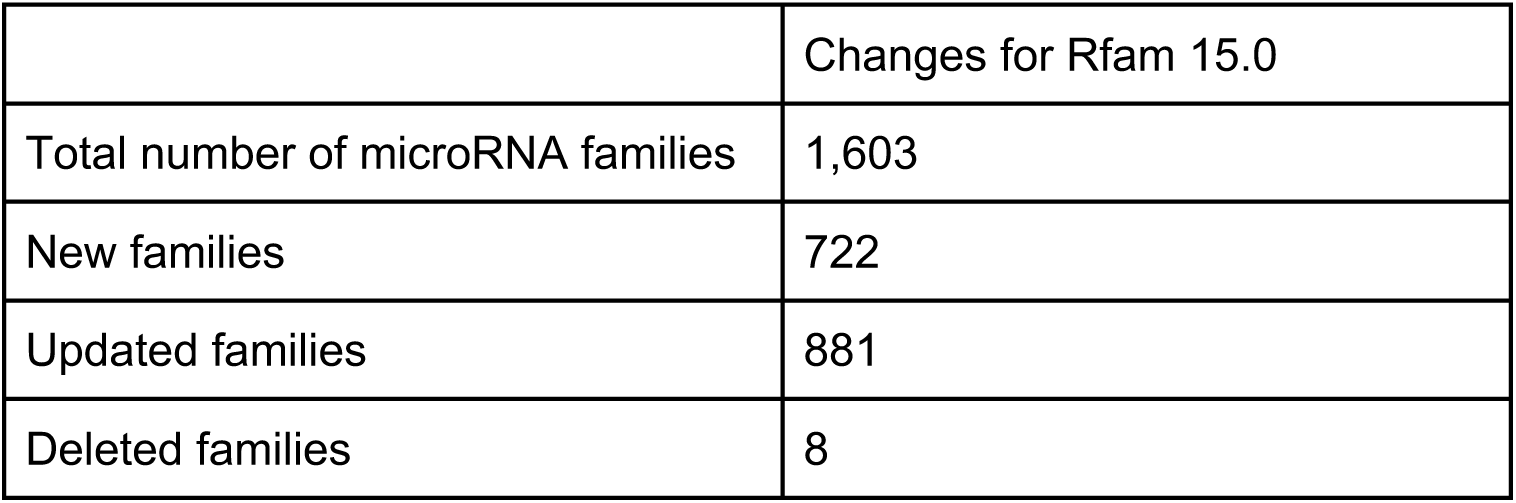
A summary of the changes in Rfam microRNA families since last publication (16) with release 14.3.

## New families

While the primary focus of Rfam has been to complete the microRNA project, and improve existing families with 3D structures, we have continued to create new families. Since Rfam 14.3 (16), Rfam has created 16 new non-microRNA or viral RNA families. The families cover a range of phylogenetic and functional types. A few examples include Bacteroidales small SRP (RF04183), the signal recognition particle RNA of Bacteroidetes (22), Hairpin-meta1, a virus-like ribozyme reported in RNA satellites of plant viruses (RF04190) (23), Hovlinc ribozyme (RF04188) (24), a newly discovered type of self-cleaving ribozymes found in human and other hominids and RF04222 PLRV xrRNA, a exoribonuclease-resistant RNA detected in Flavivirus Potato virus (25), RF04247 bZIP family which is a non canonical Hac1/Xbp1 intron found in *Metazoa* (26). Shown in Figure 4 are the secondary structures of selected families.

**Figure 4:**
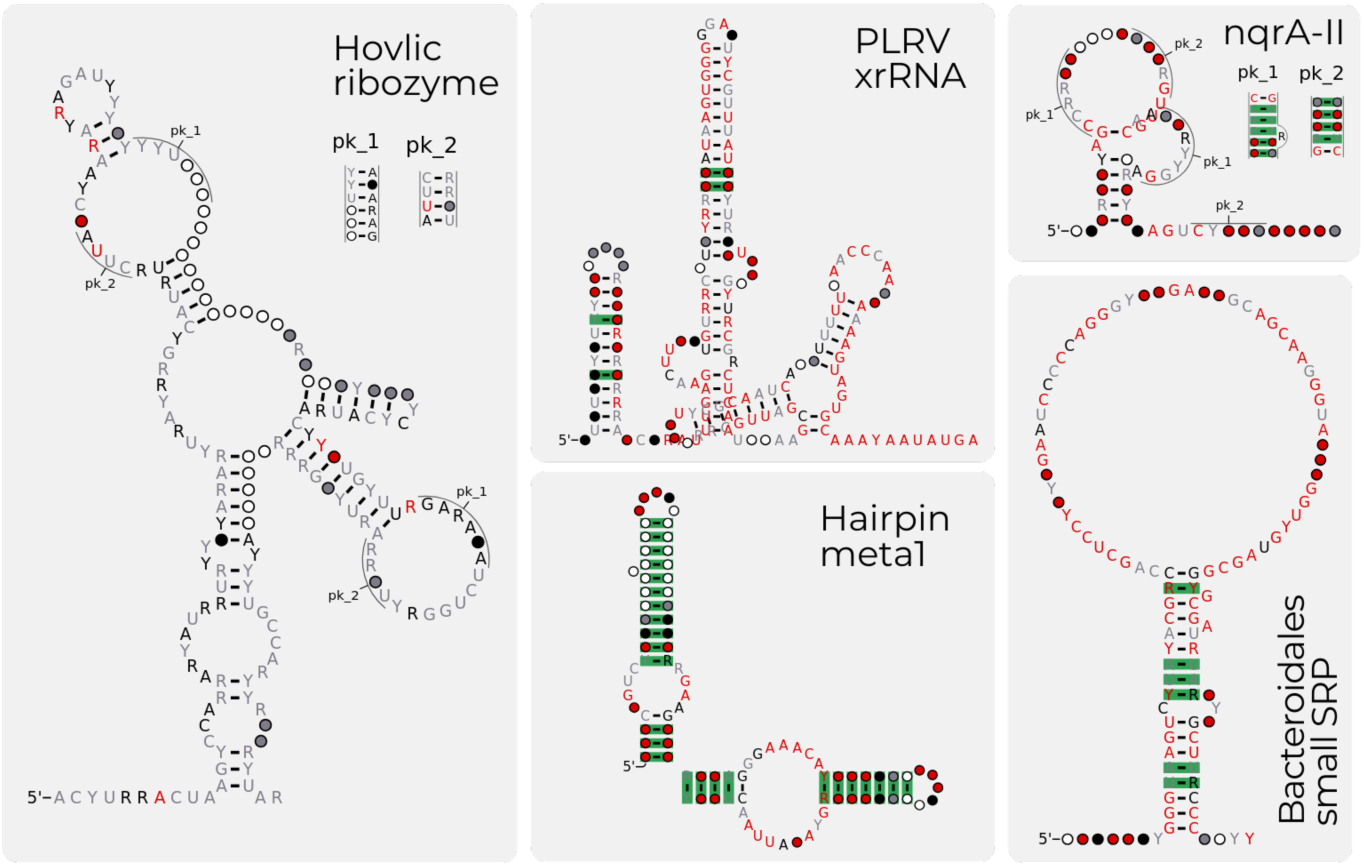
A selection of the new families for Rfam 15.0. The five families shown Hovlinc ribozyme (RF04188), the PLRV xrRNA (RF04222), the hairpin meta ribozyme (RF04190), nqrA-II ncRNA motif (RF04310) and Bacteriodales small SRP (RF04183) are several of the new families created for Rfam 15.0.

These families are curated through a combination of literature reviews and community submissions. We actively encourage groups with candidate RNAs to contribute their alignments and secondary structure data to Rfam. However, it is important to note that Rfam requires all sequences in SEED alignments to be traceable to public databases such as GenBank (27) or RNAcentral (21). With the increasing ease of sequencing, many laboratories maintain private sequence databases for constructing alignments. As a result, many of these new alignments cannot be incorporated into Rfam due to the lack of traceable public records. We strongly recommend that the scientific community submit their sequences to publicly accessible databases, ensuring they are available for reuse and analysis by the broader research community. Furthermore, when publishing alignments, using public accessions will facilitate faster and more efficient integration into resources like Rfam.

## Other improvements

### Updating FULL alignments

Rfam provides several data types for interested users to download from our FTP site. These include the SEED alignments, CMs and FULL sequences. Older versions of Rfam included FULL alignments, which are an alignment of all matching sequences to the CM. Previously, these were no longer produced because of technical limitations, however, with improvements to infernal and growth in available compute power it is now possible to create these alignments again. We now produce full alignments for all Rfam families. These are available for each model in the ‘full_alignments’ section on the FTP site (https://ftp.ebi.ac.uk/pub/databases/Rfam/CURRENT/full_alignments).

### APICURON integration

APICURON (https://apicuron.org/) is a database to track and credit biocurators for the work they do (28). The database allows for databases to credit curators for each activity they perform and provides overall statistics for each database including Rfam. In Rfam, it is common for a curator to perform many small updates and fixes to families to produce a release. Currently, this work goes unacknowledged except during publication, which are generally separated by several years. By integrating with APICURON we are able to display all the changes curators perform as part of their duties on a more regular schedule. Additionally, APICURON updates ORCID records (https://orcid.org/) to help credit curators for their activities. We update our APICURON records with each release and our APICURON page is available at: https://apicuron.org/databases/rfam. APICURON requires that each resource track the activities of each curator, unfortunately some historical Rfam activity does not have the required specificity, leading to some older curators lacking attributions. This is being worked on and we hope to properly credit all curators for their contributions to Rfam.

### Website improvements

The Rfam website (http://rfam.org) has undergone continuous development since our last publication. We have added several new features, such as dedicated landing pages for each project, e.g. viral families can easily be browsed at http://rfam.org/viruses. These pages make it simpler for users interested in a particular project, e.g. the 3D structure alignments, to see our progress and fetch all data related to the project.

We have recently updated some terminology used for riboswitch families in Rfam. We have found that some users are confused that Rfam riboswitch families generally include just the aptamer domains. This is because aptamers align well, while it is difficult to build an alignment of a whole riboswitch. We have added a short note to the header of each Rfam riboswitch page to indicate that this family is only the aptamer domain and a short explanation of what that means. Additionally, the descriptions of families now include the term ‘aptamer’ to clarify this point. We have not updated the identifiers or accessions of any families.

Finally, we have integrated the RNAcentral LitScan widget from RNAcentral. RNAcentral has developed a pipeline, LitScan, which identifies and extracts mentions of any non-coding RNA in all open access literature (29). The result of this pipeline can be reused by embedding a simple HTML widget. The widget allows users to browse and search the related literature in a convenient way. We have embedded this widget on all Rfam family pages under the new ‘Publications’ tab. The results are updated with each RNAcentral release, roughly every 4 months. An example of the widget for the Glutamine riboswitch family page is shown in Figure 5.

**Figure 5:**
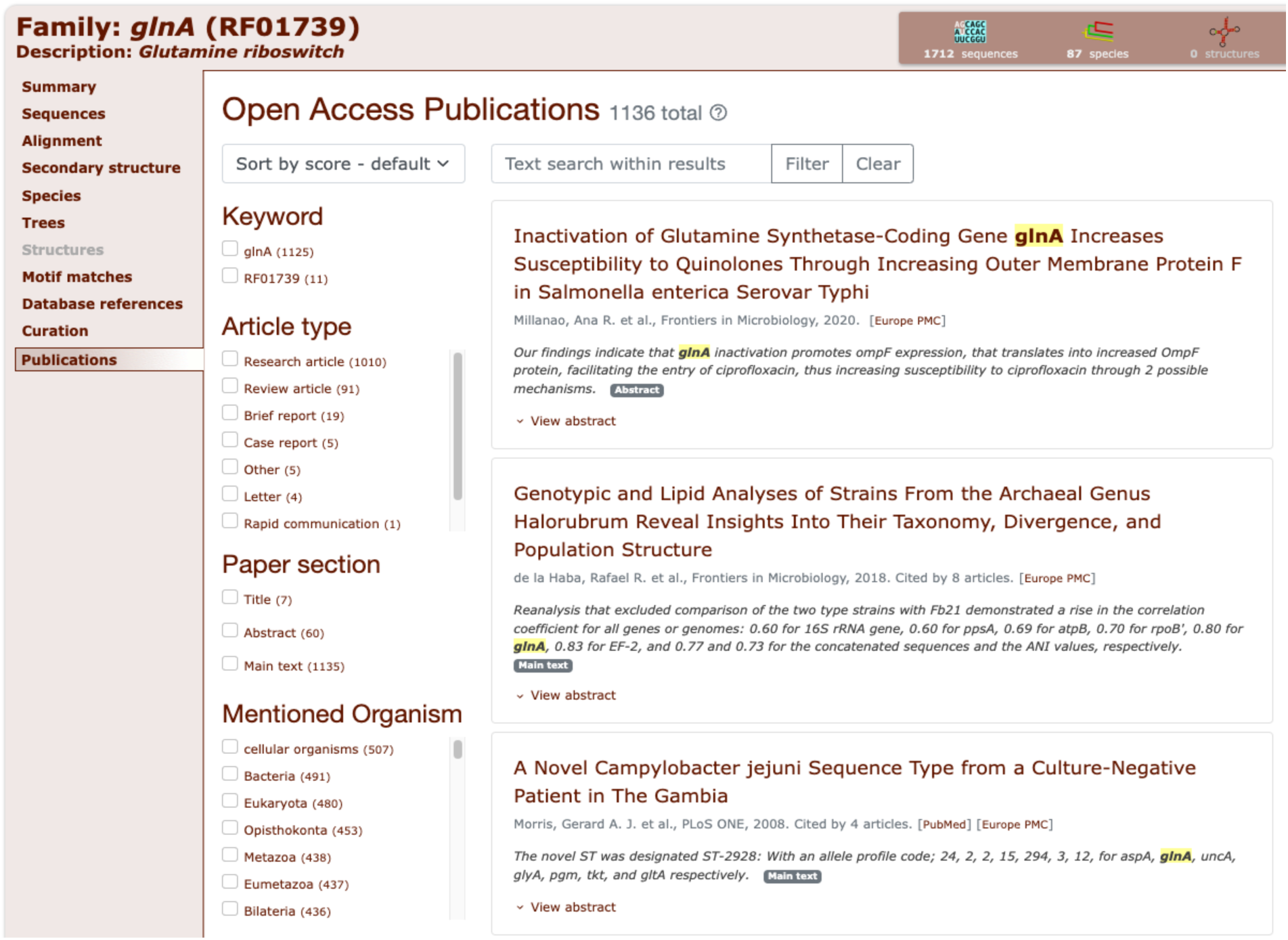
The LitScan widget embedded in the Glutamine riboswitch family page. This widget shows users which open access articles mention the family. Families are searched using their Rfam ID and Rfam accession in all open access literature. It allows users to search the matched articles by several types of metadata such as the article type (research, review, report, etc), the section of the paper which mentions the family and organisms mentioned in the paper. Additionally, users can perform a text search within the sentences which mention the Rfam family.

## Conclusions

After more than 20 years of work (1), Rfam has grown to over 4, 000 families and continues to be a key resource in RNA science. We continue to provide a large centralised collection of ncRNA families, which are used in many ways. Rfam was originally created to annotate genomes, but we have grown far beyond that use case. We would like to take this opportunity to thank everyone who has worked on or collaborated with Rfam over the past 20 years. Anyone interested in learning about the history and changes in Rfam since its inception are encouraged to visit https://rfam.org/rfam20 to find interviews with many former and current staff and users.

Since our last publication we have focused on completing our synchronisation with miRBase and improving existing families using R-scape and 3D structures. This has led to a large increase in the number and quality of families. Rfam plans to stay synchronised with miRBase and complete the 3D structure improvements by connecting each family to at least one structure where possible. Outside of those projects, we will continue to import viral RNAs and return to capturing more novel families discovered by the community. Finally, we will explore ways to expand Rfamseq to better improve our coverage of the phylogenetic space.

It is essential to note that there has been a major shift in bioinformatics with the publication of AlphaFold and its effect on protein structure prediction. We expect that as the field of RNA structure prediction matures, Rfam will continue to play a key role as a source of ground truth. In order to best serve this new use case, we will begin exploring ways to grow faster than before, while maintaining the high standards the community has come to expect of Rfam alignments. This will be essential to providing the test and training sets required for deep learning based approaches.

Finally, we encourage any interested community members to reach out for collaboration or to provide data. As discussed here, much of our major work is carried out with interested community members and can lead to new directions for the resource. We invite new data submissions and feedback at https://docs.rfam.org/page/contact-us.html.

## Data availability

All Rfam data are released under the Creative Commons Zero (CC0) licence at https://rfam.org. The data can be accessed via an API, a public MySQL database, and the FTP archive. The Rfam documentation (https://docs.rfam.org) and (30) contain detailed instructions. All code is available on GitHub under the Apache 2.0 licence at https://github.com/Rfam.

## Acknowledgements

The authors would like to thank the community for its excellent feedback and the organisers of the Benasque 2022 and 2024 meetings where several collaborations described in this paper were established. We would also like to thank the broader RNA community community for their contributions, including providing alignments and feedback for families. In particular we thank Lars Barquist, Fei Qi, Christina Weinberg, Zasha Weinberg and Quentin Vincens who kindly shared their data with us and worked with us in the creation of new families.

---

**Table S1:** The mapping between Rfam accessions, ids and miRBase family accessions.

| Rfam<br>Accession | Rfam ID | miRBase Family<br>Accession |
| --- | --- | --- |
| RF00027 | let-7 | MIPF0000002 |
| RF00047 | mir-2 | MIPF0000049 |
| RF00051 | mir-17 | MIPF0000001 |
| RF00052 | lin-4 | MIPF0000303 |
| RF00053 | mir-7 | MIPF0000022 |
| RF00073 | mir-156 | MIPF0000008 |
| RF00074 | mir-29 | MIPF0000009 |
| RF00075 | mir-166 | MIPF0000004 |
| RF00076 | mir-181 | MIPF0000007 |
| RF00103 | mir-1 | MIPF0000038 |
| RF00104 | mir-10 | MIPF0000033 |
| RF00104 | mir-10 | MIPF0000025 |
| RF00104 | mir-10 | MIPF0000268 |
| RF00104 | mir-10 | MIPF0000271 |
| RF00129 | mir-103 | MIPF0000024 |
| RF00130 | mir-192 | MIPF0000063 |
| RF00131 | mir-30 | MIPF0000005 |
| RF00143 | mir-6 | MIPF0000119 |
| RF00144 | mir-199 | MIPF0000040 |
| RF00237 | mir-9 | MIPF0000014 |
| RF00239 | mir-124 | MIPF0000021 |
| RF00241 | mir-8 | MIPF0000019 |
| RF00244 | mir-26 | MIPF0000043 |
| RF00245 | mir-19 | MIPF0000011 |
| RF00246 | mir-135 | MIPF0000028 |
| RF00247 | mir-160 | MIPF0000032 |
| RF00251 | mir-219 | MIPF0000044 |
| RF00253 | mir-101 | MIPF0000046 |
| RF00254 | mir-16 | MIPF0000006 |
| RF00255 | mir-218 | MIPF0000026 |
| RF00256 | mir-196 | MIPF0000031 |
| RF00257 | mir-194 | MIPF0000055 |
| RF00258 | mir-130 | MIPF0000034 |
| RF00363 | mir-BART1 | MIPF0000325 |
| RF00365 | mir-BHRF1-1 | MIPF0000331 |
| RF00366 | mir-BHRF1-2 | MIPF0000332 |
| RF00446 | mir-133 | MIPF0000029 |
| RF00455 | mir-15 | MIPF0000006 |
| RF00637 | mir-276 | MIPF0000124 |
| RF00638 | MIR159 | MIPF0000010 |
| RF00639 | mir-515 | MIPF0000020 |
| RF00640 | MIR167 | MIPF0000023 |
| RF00641 | mir-154 | MIPF0000018 |
| RF00642 | mir-23 | MIPF0000027 |
| RF00643 | MIR171_1 | MIPF0000030 |
| RF00644 | mir-27 | MIPF0000036 |
| RF00645 | MIR169_2 | MIPF0000037 |
| RF00646 | mir-204 | MIPF0000042 |
| RF00647 | MIR164 | MIPF0001794 |
| RF00648 | MIR396 | MIPF0000047 |
| RF00649 | mir-128 | MIPF0000048 |
| RF00650 | mir-153 | MIPF0000050 |
| RF00651 | mir-221 | MIPF0000051 |
| RF00652 | MIR478 | MIPF0000052 |
| RF00653 | mir-22 | MIPF0000053 |
| RF00654 | mir-216 | MIPF0000054 |
| RF00655 | mir-28 | MIPF0000057 |
| RF00656 | mir-205 | MIPF0000058 |
| RF00657 | mir-184 | MIPF0000059 |
| RF00658 | mir-21 | MIPF0000060 |
| RF00659 | mir-365 | MIPF0000061 |
| RF00660 | mir-214 | MIPF0000062 |
| RF00661 | mir-31 | MIPF0000064 |
| RF00662 | mir-132 | MIPF0000065 |
| RF00663 | mir-183 | MIPF0000066 |
| RF00664 | mir-223 | MIPF0000067 |
| RF00665 | mir-290 | MIPF0000068 |
| RF00666 | mir-32 | MIPF0000069 |
| RF00667 | mir-33 | MIPF0000070 |
| RF00668 | mir-302 | MIPF0000071 |
| RF00669 | mir-96 | MIPF0000072 |
| RF00670 | mir-105 | MIPF0000074 |
| RF00671 | mir-138 | MIPF0000075 |
| RF00672 | mir-190 | MIPF0000076 |
| RF00673 | mir-217 | MIPF0000077 |
| RF00674 | mir-187 | MIPF0000078 |
| RF00675 | mir-145 | MIPF0000079 |
| RF00676 | mir-127 | MIPF0000080 |
| RF00677 | MIR168 | MIPF0000081 |
| RF00678 | mir-140 | MIPF0000085 |
| RF00679 | mir-210 | MIPF0000086 |
| RF00680 | mir-224 | MIPF0000088 |
| RF00681 | mir-198 | MIPF0000090 |
| RF00682 | mir-144 | MIPF0000093 |
| RF00683 | mir-143 | MIPF0000094 |
| RF00684 | mir-122 | MIPF0000095 |
| RF00685 | mir-36 | MIPF0000096 |
| RF00686 | mir-338 | MIPF0000097 |
| RF00687 | mir-136 | MIPF0000099 |
| RF00688 | MIR394 | MIPF0000100 |
| RF00689 | MIR390 | MIPF0000101 |
| RF00690 | MIR408 | MIPF0000102 |
| RF00691 | mir-146 | MIPF0000103 |
| RF00692 | MIR171_2 | MIPF0000104 |
| RF00693 | mir-147 | MIPF0000105 |
| RF00694 | mir-137 | MIPF0000106 |
| RF00695 | MIR398 | MIPF0000107 |
| RF00696 | mir-203 | MIPF0000108 |
| RF00697 | mir-186 | MIPF0000109 |
| RF00698 | mir-489 | MIPF0000111 |
| RF00699 | mir-134 | MIPF0000112 |
| RF00700 | mir-375 | MIPF0000114 |
| RF00701 | mir-126 | MIPF0000115 |
| RF00702 | mir-182 | MIPF0000116 |
| RF00703 | mir-139 | MIPF0000117 |
| RF00704 | MIR397 | MIPF0000120 |
| RF00705 | mir-202 | MIPF0000121 |
| RF00706 | mir-263 | MIPF0000122 |
| RF00707 | mir-197 | MIPF0000123 |
| RF00708 | mir-450 | MIPF0000128 |
| RF00709 | mir-455 | MIPF0000129 |
| RF00710 | mir-44 | MIPF0000132 |
| RF00711 | mir-449 | MIPF0000133 |
| RF00712 | mir-460 | MIPF0000134 |
| RF00713 | mir-239 | MIPF0000135 |
| RF00714 | MIR535 | MIPF0000136 |
| RF00715 | mir-383 | MIPF0000137 |
| RF00716 | mir-3 | MIPF0000140 |
| RF00717 | mir-315 | MIPF0000141 |
| RF00718 | mir-431 | MIPF0000142 |
| RF00719 | mir-326 | MIPF0000143 |
| RF00720 | mir-317 | MIPF0000144 |
| RF00721 | MIR475 | MIPF0000145 |
| RF00722 | mir-451 | MIPF0000148 |
| RF00723 | mir-448 | MIPF0000149 |
| RF00724 | mir-282 | MIPF0000150 |
| RF00725 | mir-iab-4 | MIPF0000151 |
| RF00726 | mir-87 | MIPF0000152 |
| RF00727 | bantam | MIPF0000153 |
| RF00728 | mir-81 | MIPF0000154 |
| RF00729 | mir-278 | MIPF0000155 |
| RF00730 | mir-277 | MIPF0000156 |
| RF00731 | mir-155 | MIPF0000157 |
| RF00732 | mir-305 | MIPF0000158 |
| RF00733 | mir-296 | MIPF0000159 |
| RF00734 | mir-52 | MIPF0000160 |
| RF00735 | mir-367 | MIPF0000162 |
| RF00736 | mir-320 | MIPF0000163 |
| RF00737 | mir-322 | MIPF0000164 |
| RF00739 | MIR476 | MIPF0000166 |
| RF00740 | mir-370 | MIPF0000167 |
| RF00741 | mir-378 | MIPF0000168 |
| RF00742 | MIR162_1 | MIPF0000127 |
| RF00743 | mir-308 | MIPF0000171 |
| RF00744 | mir-361 | MIPF0000172 |
| RF00745 | mir-499 | MIPF0000173 |
| RF00746 | mir-454 | MIPF0000174 |
| RF00747 | mir-283 | MIPF0000175 |
| RF00748 | mir-433 | MIPF0000177 |
| RF00749 | mir-208 | MIPF0000178 |
| RF00750 | mir-458 | MIPF0000179 |
| RF00751 | mir-12 | MIPF0000181 |
| RF00752 | mir-14 | MIPF0000182 |
| RF00753 | mir-503 | MIPF0000183 |
| RF00754 | mir-279 | MIPF0000184 |
| RF00755 | mir-542 | MIPF0000185 |
| RF00756 | mir-299 | MIPF0000186 |
| RF00757 | mir-275 | MIPF0000187 |
| RF00758 | mir-346 | MIPF0000188 |
| RF00760 | mir-342 | MIPF0000190 |
| RF00761 | mir-340 | MIPF0000191 |
| RF00762 | mir-412 | MIPF0000192 |
| RF00763 | mir-339 | MIPF0000193 |
| RF00764 | mir-191 | MIPF0000194 |
| RF00765 | mir-337 | MIPF0000195 |
| RF00766 | mir-335 | MIPF0000196 |
| RF00767 | mir-150 | MIPF0000197 |
| RF00768 | MIR405 | MIPF0000198 |
| RF00769 | mir-331 | MIPF0000199 |
| RF00770 | mir-330 | MIPF0000200 |
| RF00771 | mir-185 | MIPF0000202 |
| RF00772 | mir-328 | MIPF0000203 |
| RF00773 | mir-298 | MIPF0000206 |
| RF00774 | mir-360 | MIPF0000210 |
| RF00775 | mir-432 | MIPF0000211 |
| RF00776 | mir-540 | MIPF0000212 |
| RF00777 | mir-541 | MIPF0000213 |
| RF00778 | MIR473 | MIPF0000214 |
| RF00779 | MIR474 | MIPF0000215 |
| RF00780 | MIR477 | MIPF0000216 |
| RF00781 | mir-505 | MIPF0000217 |
| RF00782 | MIR480 | MIPF0000218 |
| RF00783 | mir-484 | MIPF0000219 |
| RF00784 | mir-486 | MIPF0000220 |
| RF00785 | mir-90 | MIPF0000221 |
| RF00786 | mir-289 | MIPF0000222 |
| RF00787 | mir-288 | MIPF0000223 |
| RF00788 | mir-287 | MIPF0000224 |
| RF00789 | mir-286 | MIPF0000225 |
| RF00790 | mir-358 | MIPF0000226 |
| RF00791 | mir-357 | MIPF0000227 |
| RF00792 | mir-490 | MIPF0000229 |
| RF00793 | mir-497 | MIPF0000231 |
| RF00794 | mir-42 | MIPF0000233 |
| RF00795 | mir-43 | MIPF0000234 |
| RF00796 | mir-48 | MIPF0000235 |
| RF00797 | mir-355 | MIPF0000236 |
| RF00798 | mir-49 | MIPF0000237 |
| RF00799 | mir-354 | MIPF0000238 |
| RF00801 | mir-280 | MIPF0000240 |
| RF00802 | mir-207 | MIPF0000241 |
| RF00803 | mir-425 | MIPF0000242 |
| RF00804 | mir-240 | MIPF0000243 |
| RF00805 | mir-351 | MIPF0000244 |
| RF00806 | mir-350 | MIPF0000245 |
| RF00807 | mir-314 | MIPF0000246 |
| RF00808 | mir-86 | MIPF0000247 |
| RF00809 | mir-241 | MIPF0000248 |
| RF00810 | mir-85 | MIPF0000249 |
| RF00811 | mir-84 | MIPF0000250 |
| RF00812 | mir-83 | MIPF0000251 |
| RF00813 | mir-11 | MIPF0000252 |
| RF00814 | mir-316 | MIPF0000254 |
| RF00815 | mir-244 | MIPF0000256 |
| RF00816 | mir-245 | MIPF0000257 |
| RF00817 | mir-80 | MIPF0000258 |
| RF00818 | mir-318 | MIPF0000259 |
| RF00819 | mir-246 | MIPF0000260 |
| RF00820 | mir-248 | MIPF0000262 |
| RF00821 | mir-249 | MIPF0000263 |
| RF00822 | mir-274 | MIPF0000264 |
| RF00823 | lsy-6 | MIPF0000265 |
| RF00824 | mir-50 | MIPF0000266 |
| RF00825 | mir-344 | MIPF0000267 |
| RF00826 | mir-55 | MIPF0000270 |
| RF00827 | mir-77 | MIPF0000272 |
| RF00828 | mir-75 | MIPF0000273 |
| RF00829 | mir-149 | MIPF0000274 |
| RF00830 | mir-74 | MIPF0000275 |
| RF00831 | mir-73 | MIPF0000276 |
| RF00832 | mir-71 | MIPF0000278 |
| RF00833 | mir-70 | MIPF0000279 |
| RF00834 | mir-268 | MIPF0000280 |
| RF00835 | mir-58 | MIPF0000282 |
| RF00836 | mir-250 | MIPF0000283 |
| RF00837 | mir-251 | MIPF0000284 |
| RF00838 | mir-252 | MIPF0000285 |
| RF00839 | mir-452 | MIPF0000287 |
| RF00840 | mir-374 | MIPF0000288 |
| RF00841 | mir-384 | MIPF0000289 |
| RF00842 | MIR403 | MIPF0000290 |
| RF00843 | mir-228 | MIPF0000292 |
| RF00844 | mir-67 | MIPF0000293 |
| RF00845 | MIR158 | MIPF0000294 |
| RF00846 | mir-64 | MIPF0000295 |
| RF00847 | mir-62 | MIPF0000296 |
| RF00848 | mir-61 | MIPF0000297 |
| RF00849 | mir-60 | MIPF0000298 |
| RF00850 | mir-259 | MIPF0000299 |
| RF00851 | mir-230 | MIPF0000306 |
| RF00852 | mir-231 | MIPF0000307 |
| RF00853 | mir-304 | MIPF0000308 |
| RF00854 | mir-5 | MIPF0000309 |
| RF00855 | mir-254 | MIPF0000310 |
| RF00856 | mir-232 | MIPF0000311 |
| RF00857 | mir-233 | MIPF0000312 |
| RF00858 | mir-306 | MIPF0000313 |
| RF00859 | mir-234 | MIPF0000315 |
| RF00861 | mir-488 | MIPF0000318 |
| RF00862 | mir-491 | MIPF0000319 |
| RF00863 | mir-BART17 | MIPF0000321 |
| RF00864 | mir-BART20 | MIPF0000322 |
| RF00865 | MIR169_5 | MIPF0000323 |
| RF00866 | mir-BART3 | MIPF0000324 |
| RF00867 | mir-BART5 | MIPF0000326 |
| RF00868 | mir-BART15 | MIPF0000327 |
| RF00869 | mir-BART7 | MIPF0000328 |
| RF00870 | mir-423 | MIPF0000329 |
| RF00872 | mir-652 | MIPF0000333 |
| RF00873 | mir-550 | MIPF0000334 |
| RF00874 | mir-BART12 | MIPF0000335 |
| RF00875 | mir-692 | MIPF0000336 |
| RF00876 | mir-684 | MIPF0000337 |
| RF00877 | mir-592 | MIPF0000340 |
| RF00878 | mir-456 | MIPF0000341 |
| RF00879 | mir-615 | MIPF0000342 |
| RF00883 | MIR820 | MIPF0000348 |
| RF00884 | MIR815 | MIPF0000349 |
| RF00885 | MIR821 | MIPF0000350 |
| RF00886 | MIR807 | MIPF0000352 |
| RF00887 | mir-802 | MIPF0000353 |
| RF00888 | mir-770 | MIPF0000355 |
| RF00890 | mir-668 | MIPF0000357 |
| RF00891 | mir-671 | MIPF0000358 |
| RF00892 | mir-551 | MIPF0000360 |
| RF00893 | MIR854 | MIPF0000361 |
| RF00894 | mir-790 | MIPF0000362 |
| RF00895 | mir-786 | MIPF0000363 |
| RF00896 | mir-787 | MIPF0000364 |
| RF00897 | mir-675 | MIPF0000365 |
| RF00898 | mir-242 | MIPF0000366 |
| RF00899 | mir-235 | MIPF0000367 |
| RF00900 | mir-255 | MIPF0000368 |
| RF00901 | MIR845_1 | MIPF0000369 |
| RF00902 | mir-791 | MIPF0000370 |
| RF00903 | mir-359 | MIPF0000371 |
| RF00904 | mir-392 | MIPF0000372 |
| RF00905 | mir-789 | MIPF0000373 |
| RF00906 | MIR1122 | MIPF0000382 |
| RF00907 | mir-941 | MIPF0000387 |
| RF00908 | MIR529 | MIPF0000388 |
| RF00909 | mir-883 | MIPF0000389 |
| RF00910 | mir-873 | MIPF0000390 |
| RF00911 | mir-672 | MIPF0000391 |
| RF00912 | mir-877 | MIPF0000392 |
| RF00914 | mir-674 | MIPF0000394 |
| RF00915 | mir-760 | MIPF0000395 |
| RF00917 | mir-708 | MIPF0000397 |
| RF00918 | mir-872 | MIPF0000399 |
| RF00919 | mir-874 | MIPF0000401 |
| RF00920 | MIR444 | MIPF0000402 |
| RF00921 | mir-665 | MIPF0000404 |
| RF00922 | mir-673 | MIPF0000405 |
| RF00925 | MIR1027 | MIPF0000414 |
| RF00926 | MIR1151 | MIPF0000415 |
| RF00927 | mir-582 | MIPF0000417 |
| RF00928 | mir-590 | MIPF0000418 |
| RF00929 | mir-574 | MIPF0000419 |
| RF00931 | mir-879 | MIPF0000421 |
| RF00932 | mir-471 | MIPF0000423 |
| RF00933 | mir-875 | MIPF0000424 |
| RF00934 | mir-463 | MIPF0000425 |
| RF00935 | mir-876 | MIPF0000430 |
| RF00936 | mir-744 | MIPF0000431 |
| RF00937 | mir-653 | MIPF0000435 |
| RF00939 | mir-504 | MIPF0000437 |
| RF00940 | mir-327 | MIPF0000438 |
| RF00941 | mir-434 | MIPF0000439 |
| RF00942 | mir-1224 | MIPF0000440 |
| RF00943 | MIR824 | MIPF0000442 |
| RF00945 | mir-1226 | MIPF0000444 |
| RF00946 | mir-1225 | MIPF0000445 |
| RF00947 | mir-929 | MIPF0000447 |
| RF00948 | mir-996 | MIPF0000449 |
| RF00949 | mir-983 | MIPF0000450 |
| RF00950 | mir-927 | MIPF0000452 |
| RF00951 | mir-1302 | MIPF0000456 |
| RF00952 | mir-650 | MIPF0000457 |
| RF00953 | mir-1497 | MIPF0000458 |
| RF00954 | MIR1446 | MIPF0000459 |
| RF00955 | mir-1829 | MIPF0000460 |
| RF00956 | MIR1444 | MIPF0000461 |
| RF00957 | mir-663 | MIPF0000462 |
| RF00958 | mir-498 | MIPF0000463 |
| RF00959 | mir-612 | MIPF0000464 |
| RF00960 | mir-661 | MIPF0000465 |
| RF00961 | mir-581 | MIPF0000466 |
| RF00962 | mir-586 | MIPF0000467 |
| RF00963 | mir-642 | MIPF0000468 |
| RF00964 | mir-938 | MIPF0000469 |
| RF00965 | mir-549 | MIPF0000470 |
| RF00966 | mir-676 | MIPF0000471 |
| RF00967 | mir-281 | MIPF0000472 |
| RF00968 | mir-626 | MIPF0000474 |
| RF00969 | mir-556 | MIPF0000475 |
| RF00970 | mir-648 | MIPF0000476 |
| RF00971 | mir-578 | MIPF0000477 |
| RF00972 | mir-651 | MIPF0000478 |
| RF00973 | mir-597 | MIPF0000479 |
| RF00974 | mir-607 | MIPF0000480 |
| RF00975 | MIR845_2 | MIPF0000481 |
| RF00976 | mir-583 | MIPF0000482 |
| RF00977 | mir-600 | MIPF0000484 |
| RF00978 | mir-638 | MIPF0000486 |
| RF00979 | mir-553 | MIPF0000487 |
| RF00980 | mir-643 | MIPF0000488 |
| RF00981 | mir-939 | MIPF0000490 |
| RF00983 | mir-662 | MIPF0000492 |
| RF00984 | mir-576 | MIPF0000493 |
| RF00985 | mir-640 | MIPF0000494 |
| RF00986 | mir-920 | MIPF0000495 |
| RF00987 | mir-589 | MIPF0000496 |
| RF00988 | mir-657 | MIPF0000498 |
| RF00989 | mir-492 | MIPF0000499 |
| RF00990 | mir-552 | MIPF0000501 |
| RF00991 | mir-599 | MIPF0000502 |
| RF00992 | mir-593 | MIPF0000503 |
| RF00993 | mir-1473 | MIPF0000504 |
| RF00994 | mir-1255 | MIPF0000506 |
| RF00995 | mir-616 | MIPF0000507 |
| RF00996 | mir-631 | MIPF0000508 |
| RF00997 | mir-942 | MIPF0000511 |
| RF00998 | mir-562 | MIPF0000512 |
| RF00999 | mir-924 | MIPF0000513 |
| RF01000 | mir-580 | MIPF0000515 |
| RF01001 | mir-609 | MIPF0000516 |
| RF01002 | mir-936 | MIPF0000517 |
| RF01003 | mir-563 | MIPF0000519 |
| RF01004 | mir-557 | MIPF0000520 |
| RF01005 | MIR530 | MIPF0000521 |
| RF01006 | mir-601 | MIPF0000522 |
| RF01007 | mir-624 | MIPF0000523 |
| RF01008 | mir-636 | MIPF0000524 |
| RF01009 | mir-M7 | MIPF0000526 |
| RF01010 | mir-632 | MIPF0000527 |
| RF01011 | mir-605 | MIPF0000528 |
| RF01012 | mir-628 | MIPF0000529 |
| RF01013 | mir-577 | MIPF0000530 |
| RF01014 | mir-1306 | MIPF0000531 |
| RF01015 | mir-885 | MIPF0000532 |
| RF01016 | mir-584 | MIPF0000533 |
| RF01018 | mir-569 | MIPF0000535 |
| RF01019 | mir-922 | MIPF0000536 |
| RF01020 | mir-572 | MIPF0000537 |
| RF01021 | mir-558 | MIPF0000538 |
| RF01022 | mir-611 | MIPF0000539 |
| RF01023 | mir-940 | MIPF0000540 |
| RF01024 | mir-944 | MIPF0000541 |
| RF01025 | mir-934 | MIPF0000542 |
| RF01026 | MIR828 | MIPF0000544 |
| RF01027 | mir-765 | MIPF0000545 |
| RF01028 | mir-633 | MIPF0000546 |
| RF01029 | mir-649 | MIPF0000547 |
| RF01030 | mir-422 | MIPF0000548 |
| RF01031 | mir-639 | MIPF0000549 |
| RF01032 | mir-554 | MIPF0000551 |
| RF01033 | mir-767 | MIPF0000552 |
| RF01034 | mir-618 | MIPF0000553 |
| RF01035 | mir-887 | MIPF0000554 |
| RF01036 | mir-567 | MIPF0000555 |
| RF01037 | mir-644 | MIPF0000556 |
| RF01038 | mir-1307 | MIPF0000558 |
| RF01039 | mir-937 | MIPF0000560 |
| RF01040 | mir-573 | MIPF0000561 |
| RF01041 | mir-604 | MIPF0000562 |
| RF01042 | mir-891 | MIPF0000420 |
| RF01043 | MIR1023 | MIPF0000396 |
| RF01044 | mir-345 | MIPF0000189 |
| RF01045 | mir-544 | MIPF0000436 |
| RF01059 | mir-598 | MIPF0000393 |
| RF01061 | mir-548 | MIPF0000317 |
| RF01064 | mir-253 | MIPF0000286 |
| RF01117 | ciona-mir-92 | MIPF0000509 |
| RF01314 | mir-1227 | MIPF0000454 |
| RF01413 | miR-430 | MIPF0000003 |
| RF01895 | mir-193 | MIPF0000082 |
| RF01896 | mir-142 | MIPF0000084 |
| RF01897 | mir-188 | MIPF0000113 |
| RF01898 | mir-363 | MIPF0000138 |
| RF01899 | mir-2241 | MIPF0000821 |
| RF01900 | mir-2024 | MIPF0000697 |
| RF01901 | mir-284 | MIPF0000228 |
| RF01902 | MIR439 | MIPF0000092 |
| RF01903 | mir-500 | MIPF0000139 |
| RF01910 | mir-511 | MIPF0000130 |
| RF01911 | MIR2118 | MIPF0000745 |
| RF01912 | mir-2807 | MIPF0000826 |
| RF01913 | mir-2778 | MIPF0000827 |
| RF01914 | mir-932 | MIPF0000588 |
| RF01915 | mir-2238 | MIPF0000822 |
| RF01916 | mir-988 | MIPF0000724 |
| RF01917 | MIR2587 | MIPF0000835 |
| RF01918 | mir-1249 | MIPF0000667 |
| RF01919 | mir-1419 | MIPF0000570 |
| RF01920 | mir-764 | MIPF0000707 |
| RF01921 | mir-1296 | MIPF0000649 |
| RF01922 | mir-654 | MIPF0000409 |
| RF01923 | mir-711 | MIPF0000748 |
| RF01924 | mir-2774 | MIPF0000829 |
| RF01925 | MIR1428 | MIPF0000575 |
| RF01926 | mir-981 | MIPF0000710 |
| RF01927 | MIR1222 | MIPF0000410 |
| RF01936 | mir-63 | MIPF0000823 |
| RF01937 | mir-2780 | MIPF0000828 |
| RF01938 | mir-1251 | MIPF0000621 |
| RF01939 | mir-761 | MIPF0000709 |
| RF01940 | hvt-mir-H | MIPF0000783 |
| RF01941 | MIR1223 | MIPF0000383 |
| RF01943 | mir-999 | MIPF0000852 |
| RF01944 | mir-2518 | MIPF0000858 |
| RF01945 | mir-1388 | MIPF0000805 |
| RF01996 | mir-995 | MIPF0000895 |
| RF01997 | mir-969 | MIPF0000879 |
| RF02000 | MIR1846 | MIPF0000577 |
| RF02006 | mir-1253 | MIPF0000611 |
| RF02007 | mir-1237 | MIPF0000585 |
| RF02008 | mir-621 | MIPF0000631 |
| RF02009 | mir-987 | MIPF0000859 |
| RF02010 | mir-3180 | MIPF0000894 |
| RF02011 | mir-575 | MIPF0000586 |
| RF02014 | mir-1178 | MIPF0000601 |
| RF02015 | mir-1287 | MIPF0000725 |
| RF02016 | mir-1183 | MIPF0000668 |
| RF02017 | mir-1912 | MIPF0000768 |
| RF02018 | mir-1207 | MIPF0000596 |
| RF02019 | mir-1265 | MIPF0000688 |
| RF02020 | mir-25 | MIPF0000013 |
| RF02021 | mir-3179 | MIPF0000900 |
| RF02022 | mir-1275 | MIPF0000674 |
| RF02023 | mir-1208 | MIPF0000628 |
| RF02024 | mir-1180 | MIPF0000789 |
| RF02025 | mir-3017 | MIPF0001107 |
| RF02026 | mir-2833 | MIPF0000886 |
| RF02027 | MIR2907 | MIPF0000861 |
| RF02028 | mir-1827 | MIPF0000646 |
| RF02092 | mir-2970 | MIPF0001328 |
| RF02095 | mir-2985 | MIPF0001067 |
| RF02516 | mir-393 | MIPF0000083 |
| RF02518 | mir-2494 | MIPF0000975 |
| RF02520 | mir-965 | MIPF0000708 |
| RF03171 | mir-1012 | MIPF0001046 |
| RF03172 | mir-6012 | MIPF0001496 |
| RF03173 | mir-4504 | MIPF0001508 |
| RF03174 | mir-4803 | MIPF0001511 |
| RF03175 | MIR3454 | MIPF0001097 |
| RF03176 | MIR7729 | MIPF0001721 |
| RF03177 | mir-2067 | MIPF0001854 |
| RF03178 | mir-4515 | MIPF0001512 |
| RF03179 | mir-1182 | MIPF0000673 |
| RF03180 | mir-3126 | MIPF0001525 |
| RF03181 | mir-2755 | MIPF0001452 |
| RF03182 | MIR6032 | MIPF0002003 |
| RF03183 | MIR5516 | MIPF0001595 |
| RF03184 | mir-1250 | MIPF0000696 |
| RF03185 | mir-3193 | MIPF0001401 |
| RF03186 | mir-3689 | MIPF0001144 |
| RF03187 | MIR1512 | MIPF0001291 |
| RF03188 | mir-2843 | MIPF0001044 |
| RF03189 | mir-1175 | MIPF0000715 |
| RF03190 | mir-4857 | MIPF0001842 |
| RF03191 | mir-1913 | MIPF0001015 |
| RF03192 | mir-6794 | MIPF0001726 |
| RF03193 | mir-2941 | MIPF0000842 |
| RF03194 | mir-1272 | MIPF0000645 |
| RF03195 | mir-3934 | MIPF0001584 |
| RF03196 | mir-6790 | MIPF0001811 |
| RF03197 | mir-4887 | MIPF0001841 |
| RF03198 | mir-7385 | MIPF0001688 |
| RF03199 | MIR5271 | MIPF0001281 |
| RF03200 | mir-BART13 | MIPF0001051 |
| RF03201 | mir-617 | MIPF0000672 |
| RF03202 | mir-555 | MIPF0000583 |
| RF03203 | mir-3070 | MIPF0001172 |
| RF03204 | mir-4510 | MIPF0001467 |
| RF03205 | mir-1898 | MIPF0000786 |
| RF03206 | mir-3102 | MIPF0001469 |
| RF03207 | mir-3960 | MIPF0001271 |
| RF03208 | mir-1781 | MIPF0001583 |
| RF03209 | MIR9657 | MIPF0001955 |
| RF03210 | mir-7359 | MIPF0001650 |
| RF03211 | mir-666 | MIPF0000782 |
| RF03212 | mir-2788 | MIPF0001204 |
| RF03213 | mir-5885 | MIPF0001404 |
| RF03214 | mir-769 | MIPF0000727 |
| RF03215 | mir-5360 | MIPF0001723 |
| RF03216 | mir-BART10 | MIPF0000850 |
| RF03217 | mir-3167 | MIPF0001823 |
| RF03218 | mir-3015 | MIPF0001111 |
| RF03219 | mir-602 | MIPF0000767 |
| RF03220 | mir-2582 | MIPF0000870 |
| RF03221 | mir-6715 | MIPF0001708 |
| RF03222 | mir-4900 | MIPF0001224 |
| RF03223 | mir-2209 | MIPF0000787 |
| RF03224 | MIR9560 | MIPF0002116 |
| RF03225 | mir-2763 | MIPF0001437 |
| RF03226 | mir-8196 | MIPF0002057 |
| RF03227 | mir-2528 | MIPF0001059 |
| RF03228 | mir-1471 | MIPF0001069 |
| RF03229 | mir-1297 | MIPF0000584 |
| RF03230 | mir-967 | MIPF0001096 |
| RF03231 | mir-4523 | MIPF0002104 |
| RF03232 | mir-6526 | MIPF0001431 |
| RF03233 | hvt-mir-H16 | MIPF0000773 |
| RF03234 | mir-2813 | MIPF0001043 |
| RF03235 | mir-2543 | MIPF0000856 |
| RF03236 | mir-1206 | MIPF0000634 |
| RF03237 | MIR6425 | MIPF0001386 |
| RF03238 | mir-1542 | MIPF0000485 |
| RF03239 | mir-963 | MIPF0001035 |
| RF03240 | MIR5534 | MIPF0001324 |
| RF03241 | mir-1789 | MIPF0001636 |
| RF03242 | mir-1397 | MIPF0000642 |
| RF03243 | mir-2731 | MIPF0000795 |
| RF03244 | mir-4798 | MIPF0001574 |
| RF03246 | MIR1515 | MIPF0001591 |
| RF03247 | mir-1324 | MIPF0000589 |
| RF03248 | MIR9772 | MIPF0001934 |
| RF03249 | mir-2114 | MIPF0001464 |
| RF03251 | mir-4446 | MIPF0001385 |
| RF03252 | mir-678 | MIPF0000774 |
| RF03253 | mir-4654 | MIPF0001507 |
| RF03254 | mir-9460 | MIPF0001910 |
| RF03255 | mir-2188 | MIPF0000812 |
| RF03256 | mir-1286 | MIPF0000682 |
| RF03257 | mir-4427 | MIPF0001640 |
| RF03258 | mir-4 | MIPF0000305 |
| RF03259 | MIR1514 | MIPF0000694 |
| RF03260 | mir-8229 | MIPF0001992 |
| RF03261 | mir-1759 | MIPF0001565 |
| RF03262 | mir-1181 | MIPF0000665 |
| RF03263 | MIR861 | MIPF0001190 |
| RF03264 | MIR1024 | MIPF0000406 |
| RF03265 | mir-1460 | MIPF0001502 |
| RF03266 | mir-2943 | MIPF0001056 |
| RF03267 | mir-645 | MIPF0000657 |
| RF03268 | mir-1911 | MIPF0001787 |
| RF03269 | mir-5697 | MIPF0001748 |
| RF03270 | MIR3712 | MIPF0001625 |
| RF03271 | mir-1247 | MIPF0000669 |
| RF03272 | mir-2187 | MIPF0001770 |
| RF03274 | mir-1238 | MIPF0000941 |
| RF03275 | mir-2542 | MIPF0000889 |
| RF03276 | mir-2232 | MIPF0001054 |
| RF03277 | mir-5886 | MIPF0001638 |
| RF03278 | mir-980 | MIPF0000890 |
| RF03279 | mir-1193 | MIPF0000714 |
| RF03280 | mir-7583 | MIPF0001676 |
| RF03281 | mir-596 | MIPF0000808 |
| RF03283 | mir-1010 | MIPF0000866 |
| RF03284 | mir-4637 | MIPF0001586 |
| RF03285 | mir-564 | MIPF0000612 |
| RF03286 | MIR6146 | MIPF0001562 |
| RF03287 | mir-5904 | MIPF0001444 |
| RF03288 | mir-3085 | MIPF0001193 |
| RF03289 | mir-1205 | MIPF0000609 |
| RF03290 | mir-9428 | MIPF0001958 |
| RF03291 | mir-591 | MIPF0000641 |
| RF03292 | mir-1301 | MIPF0000742 |
| RF03293 | mir-4672 | MIPF0001551 |
| RF03294 | mir-H3 | MIPF0000717 |
| RF03295 | mir-7594 | MIPF0001674 |
| RF03296 | MIR8788 | MIPF0001903 |
| RF03297 | MIR1033 | MIPF0000385 |
| RF03298 | mir-4488 | MIPF0001623 |
| RF03299 | mir-1000 | MIPF0000701 |
| RF03300 | mir-3617 | MIPF0001477 |
| RF03301 | mir-3154 | MIPF0001387 |
| RF03302 | mir-6133 | MIPF0001415 |
| RF03303 | mir-973 | MIPF0001970 |
| RF03304 | mir-1744 | MIPF0001589 |
| RF03305 | mir-3338 | MIPF0001449 |
| RF03306 | mir-3923 | MIPF0001527 |
| RF03307 | mir-1825 | MIPF0000670 |
| RF03308 | MIR848 | MIPF0001140 |
| RF03309 | mir-4791 | MIPF0001529 |
| RF03310 | mir-1544 | MIPF0001645 |
| RF03311 | MIR169 | MIPF0000832 |
| RF03312 | mir-880 | MIPF0000433 |
| RF03313 | mir-1468 | MIPF0000777 |
| RF03314 | mir-1908 | MIPF0001021 |
| RF03315 | mir-723 | MIPF0001831 |
| RF03316 | mir-1263 | MIPF0000687 |
| RF03317 | mir-4874 | MIPF0001843 |
| RF03318 | mir-2208 | MIPF0000772 |
| RF03319 | mir-989 | MIPF0000885 |
| RF03320 | mir-1743 | MIPF0001544 |
| RF03321 | mir-1006 | MIPF0000873 |
| RF03322 | mir-1641 | MIPF0001533 |
| RF03323 | mir-3613 | MIPF0001411 |
| RF03324 | mir-1288 | MIPF0000660 |
| RF03325 | mir-4872 | MIPF0001205 |
| RF03326 | mir-312 | MIPF0000573 |
| RF03327 | mir-1266 | MIPF0000615 |
| RF03328 | mir-606 | MIPF0000780 |
| RF03329 | mir-4743 | MIPF0001384 |
| RF03330 | mir-750 | MIPF0000796 |
| RF03331 | mir-247 | MIPF0001019 |
| RF03332 | mir-2237 | MIPF0000883 |
| RF03333 | mir-2820 | MIPF0001034 |
| RF03334 | MIR391 | MIPF0001151 |
| RF03335 | MIR2950 | MIPF0001121 |
| RF03336 | mir-1322 | MIPF0000663 |
| RF03337 | mir-6505 | MIPF0001805 |
| RF03338 | MIR4416 | MIPF0001389 |
| RF03339 | mir-3047 | MIPF0001196 |
| RF03340 | mir-6511 | MIPF0001382 |
| RF03341 | mir-943 | MIPF0000690 |
| RF03342 | mir-1467 | MIPF0001416 |
| RF03343 | mir-3347 | MIPF0001022 |
| RF03344 | mir-5365 | MIPF0001363 |
| RF03345 | MIR3444 | MIPF0001177 |
| RF03346 | mir-881 | MIPF0000429 |
| RF03347 | mir-2816 | MIPF0000884 |
| RF03348 | mir-667 | MIPF0000794 |
| RF03349 | mir-3151 | MIPF0001394 |
| RF03350 | mir-3851 | MIPF0001201 |
| RF03351 | MIR6020 | MIPF0001610 |
| RF03352 | mir-4869 | MIPF0001829 |
| RF03353 | MIR5291 | MIPF0001293 |
| RF03354 | mir-459 | MIPF0001322 |
| RF03355 | mir-2008 | MIPF0000729 |
| RF03356 | mir-4782 | MIPF0001546 |
| RF03357 | mir-1842 | MIPF0000788 |
| RF03358 | mir-634 | MIPF0000600 |
| RF03359 | mir-1729 | MIPF0000816 |
| RF03360 | mir-3156 | MIPF0000891 |
| RF03361 | mir-3136 | MIPF0001575 |
| RF03362 | mir-9436 | MIPF0002111 |
| RF03363 | mir-6129 | MIPF0001423 |
| RF03364 | mir-427 | MIPF0000441 |
| RF03365 | mir-1236 | MIPF0000689 |
| RF03366 | mir-3122 | MIPF0001549 |
| RF03367 | mir-975 | MIPF0001038 |
| RF03368 | mir-1822 | MIPF0001009 |
| RF03369 | mir-2217 | MIPF0001912 |
| RF03370 | mir-2566 | MIPF0000880 |
| RF03371 | mir-4788 | MIPF0001573 |
| RF03372 | mir-5918 | MIPF0001395 |
| RF03373 | mir-3170 | MIPF0001637 |
| RF03374 | mir-2001 | MIPF0000711 |
| RF03375 | MIR1094 | MIPF0000400 |
| RF03376 | mir-3075 | MIPF0001489 |
| RF03377 | mir-960 | MIPF0000897 |
| RF03378 | mir-7398 | MIPF0001643 |
| RF03379 | mir-4667 | MIPF0001390 |
| RF03380 | mir-9441 | MIPF0002092 |
| RF03381 | mir-2796 | MIPF0001057 |
| RF03382 | mir-635 | MIPF0000662 |
| RF03383 | mir-1976 | MIPF0001633 |
| RF03384 | mir-7180 | MIPF0002036 |
| RF03385 | mir-614 | MIPF0000647 |
| RF03386 | mir-1994 | MIPF0000743 |
| RF03387 | mir-1902 | MIPF0000791 |
| RF03388 | mir-9418 | MIPF0002040 |
| RF03389 | mir-4738 | MIPF0001521 |
| RF03390 | mir-2450 | MIPF0000758 |
| RF03391 | mir-588 | MIPF0000681 |
| RF03392 | mir-3943 | MIPF0001400 |
| RF03393 | mir-2849 | MIPF0000876 |
| RF03394 | mir-2574 | MIPF0001029 |
| RF03395 | mir-718 | MIPF0000721 |
| RF03396 | mir-2814 | MIPF0001070 |
| RF03397 | mir-1234 | MIPF0000593 |
| RF03398 | mir-3552 | MIPF0002093 |
| RF03399 | mir-986 | MIPF0000860 |
| RF03400 | mir-728 | MIPF0001670 |
| RF03401 | mir-1323 | MIPF0000617 |
| RF03402 | mir-957 | MIPF0000723 |
| RF03403 | mir-5366 | MIPF0001738 |
| RF03404 | mir-1298 | MIPF0000598 |
| RF03405 | mir-3200 | MIPF0001441 |
| RF03406 | mir-3099 | MIPF0001559 |
| RF03407 | MIR531 | MIPF0000607 |
| RF03408 | mir-5890 | MIPF0001378 |
| RF03409 | mir-646 | MIPF0000605 |
| RF03410 | mir-1891 | MIPF0000857 |
| RF03411 | mir-2856 | MIPF0001071 |
| RF03412 | mir-1915 | MIPF0001037 |
| RF03413 | MIR904 | MIPF0000379 |
| RF03414 | mir-670 | MIPF0000734 |
| RF03416 | mir-7383 | MIPF0001662 |
| RF03417 | mir-343 | MIPF0000411 |
| RF03418 | mir-3198 | MIPF0001216 |
| RF03419 | mir-998 | MIPF0000952 |
| RF03420 | mir-1200 | MIPF0001036 |
| RF03421 | mir-2558 | MIPF0000817 |
| RF03422 | mir-637 | MIPF0000597 |
| RF03423 | mir-962 | MIPF0001011 |
| RF03424 | mir-4856 | MIPF0001847 |
| RF03425 | mir-1292 | MIPF0000675 |
| RF03426 | mir-1294 | MIPF0000603 |
| RF03427 | mir-1005 | MIPF0000943 |
| RF03428 | mir-6134 | MIPF0001445 |
| RF03429 | mir-4451 | MIPF0001582 |
| RF03430 | MIR5294 | MIPF0001303 |
| RF03431 | mir-4657 | MIPF0002009 |
| RF03432 | mir-7331 | MIPF0001736 |
| RF03433 | mir-2576 | MIPF0001066 |
| RF03434 | mir-1246 | MIPF0000627 |
| RF03435 | mir-726 | MIPF0001344 |
| RF03436 | mir-759 | MIPF0000706 |
| RF03437 | mir-2392 | MIPF0001614 |
| RF03438 | mir-1956 | MIPF0001470 |
| RF03439 | mir-1791 | MIPF0001396 |
| RF03440 | MIR4414 | MIPF0001299 |
| RF03441 | mir-613 | MIPF0000590 |
| RF03442 | mir-4003 | MIPF0000840 |
| RF03443 | mir-2131 | MIPF0001631 |
| RF03444 | MIR5635 | MIPF0001283 |
| RF03445 | mir-7552 | MIPF0001916 |
| RF03446 | mir-2954 | MIPF0001032 |
| RF03447 | mir-1231 | MIPF0001309 |
| RF03448 | mir-6131 | MIPF0001412 |
| RF03449 | mir-6827 | MIPF0001747 |
| RF03450 | MIR1513 | MIPF0001360 |
| RF03451 | mir-3188 | MIPF0001488 |
| RF03452 | mir-3121 | MIPF0001417 |
| RF03453 | mir-1276 | MIPF0000652 |
| RF03454 | mir-1267 | MIPF0000640 |
| RF03455 | mir-9847 | MIPF0002065 |
| RF03456 | mir-724 | MIPF0001616 |
| RF03457 | MIR173 | MIPF0001175 |
| RF03458 | mir-766 | MIPF0000677 |
| RF03459 | mir-1890 | MIPF0000848 |
| RF03460 | mir-561 | MIPF0000623 |
| RF03461 | MIR842 | MIPF0001158 |
| RF03462 | mir-4677 | MIPF0001848 |
| RF03463 | mir-3036 | MIPF0001098 |
| RF03464 | mir-465 | MIPF0000384 |
| RF03465 | MIR9666 | MIPF0001930 |
| RF03466 | mir-1184 | MIPF0000592 |
| RF03467 | mir-2851 | MIPF0001014 |
| RF03468 | mir-2830 | MIPF0001052 |
| RF03469 | mir-1179 | MIPF0000632 |
| RF03470 | mir-571 | MIPF0000757 |
| RF03471 | mir-745 | MIPF0001267 |
| RF03472 | mir-623 | MIPF0000798 |
| RF03473 | mir-8813 | MIPF0002069 |
| RF03474 | mir-4766 | MIPF0001764 |
| RF03475 | mir-2387 | MIPF0001986 |
| RF03476 | mir-7360 | MIPF0001649 |
| RF03477 | MIR862_2 | MIPF0001295 |
| RF03478 | mir-630 | MIPF0000653 |
| RF03479 | mir-324 | MIPF0000165 |
| RF03480 | mir-1284 | MIPF0000604 |
| RF03481 | mir-9014 | MIPF0002031 |
| RF03482 | mir-4774 | MIPF0001578 |
| RF03483 | MIR319 | MIPF0001104 |
| RF03484 | mir-K12-10 | MIPF0000261 |
| RF03485 | mir-4000 | MIPF0000867 |
| RF03486 | mir-3940 | MIPF0001599 |
| RF03487 | mir-4423 | MIPF0001587 |
| RF03488 | mir-2366 | MIPF0001403 |
| RF03489 | mir-4796 | MIPF0001402 |
| RF03490 | mir-1426 | MIPF0000692 |
| RF03491 | mir-3146 | MIPF0001626 |
| RF03492 | mir-878 | MIPF0000434 |
| RF03493 | mir-1281 | MIPF0000695 |
| RF03494 | mir-4680 | MIPF0001988 |
| RF03495 | mir-968 | MIPF0000865 |
| RF03496 | MIR2086 | MIPF0001302 |
| RF03497 | MIR5038 | MIPF0001368 |
| RF03498 | mir-3145 | MIPF0001427 |
| RF03499 | mir-702 | MIPF0001163 |
| RF03500 | mir-1262 | MIPF0000595 |
| RF03501 | mir-3140 | MIPF0001615 |
| RF03502 | mir-1914 | MIPF0001040 |
| RF03503 | mir-547 | MIPF0000779 |
| RF03504 | mir-9461 | MIPF0002000 |
| RF03505 | mir-9128 | MIPF0002044 |
| RF03506 | mir-587 | MIPF0000563 |
| RF03507 | mir-4429 | MIPF0001474 |
| RF03508 | mir-970 | MIPF0000700 |
| RF03509 | mir-4660 | MIPF0001460 |
| RF03510 | mir-3660 | MIPF0001397 |
| RF03511 | mir-5444 | MIPF0001358 |
| RF03512 | mir-3479 | MIPF0001629 |
| RF03514 | mir-6128 | MIPF0001442 |
| RF03516 | MIR5229 | MIPF0001323 |
| RF03517 | MIR5225 | MIPF0001388 |
| RF03518 | mir-1893 | MIPF0001812 |
| RF03519 | mir-238 | MIPF0000862 |
| RF03520 | mir-3059 | MIPF0001766 |
| RF03521 | mir-23c | MIPF0001429 |
| RF03522 | mir-4850 | MIPF0002086 |
| RF03523 | mir-4703 | MIPF0001759 |
| RF03524 | mir-3174 | MIPF0001569 |
| RF03525 | mir-1343 | MIPF0001206 |
| RF03526 | mir-7177 | MIPF0002020 |
| RF03527 | mir-3072 | MIPF0001410 |
| RF03528 | mir-1264 | MIPF0000098 |
| RF03548 | mir-302_2 | MIPF0000658 |
| RF03549 | mir-15_2 | MIPF0001828 |
| RF03550 | mir-31_2 | MIPF0001803 |
| RF03552 | mir-43_2 | MIPF0001285 |
| RF03553 | mir-190_2 | MIPF0001853 |
| RF03554 | mir-92_2 | MIPF0001099 |
| RF03555 | mir-2_2 | MIPF0001838 |
| RF03556 | mir-559 | MIPF0000656 |
| RF03557 | MIR6019 | MIPF0001486 |
| RF03558 | mir-629 | MIPF0001555 |
| RF03559 | mir-8303 | MIPF0002121 |
| RF03560 | mir-3926 | MIPF0001118 |
| RF03561 | mir-3533 | MIPF0002063 |
| RF03562 | mir-2004 | MIPF0002042 |
| RF03563 | mir-BART16 | MIPF0000991 |
| RF03564 | mir-9416 | MIPF0002105 |
| RF03565 | mir-1282 | MIPF0000587 |
| RF03566 | mir-4875 | MIPF0001222 |
| RF03567 | MIR8024 | MIPF0001789 |
| RF03568 | mir-3204 | MIPF0000908 |
| RF03569 | mir-7973 | MIPF0001693 |
| RF03570 | mir-8286 | MIPF0001964 |
| RF03571 | mir-734 | MIPF0001543 |
| RF03572 | MIR2109 | MIPF0001000 |
| RF03573 | mir-1954 | MIPF0001275 |
| RF03574 | mir-4847 | MIPF0002100 |
| RF03575 | mir-rL1-14 | MIPF0000993 |
| RF03576 | mir-4435 | MIPF0001220 |
| RF03577 | mir-9193 | MIPF0002089 |
| RF03578 | mir-3910 | MIPF0001148 |
| RF03579 | mir-2058 | MIPF0001793 |
| RF03580 | mir-928 | MIPF0000950 |
| RF03581 | mir-4878 | MIPF0001827 |
| RF03582 | mir-m22-1 | MIPF0000451 |
| RF03583 | MIR781 | MIPF0001137 |
| RF03584 | mir-2290 | MIPF0001973 |
| RF03585 | mir-3938 | MIPF0001520 |
| RF03586 | mir-5132 | MIPF0002061 |
| RF03587 | mir-1202 | MIPF0000613 |
| RF03588 | mir-2319 | MIPF0000790 |
| RF03589 | mir-5583 | MIPF0001356 |
| RF03590 | mir-2009 | MIPF0001923 |
| RF03591 | mir-3117 | MIPF0001506 |
| RF03592 | mir-2567 | MIPF0000934 |
| RF03593 | mir-2331 | MIPF0001951 |
| RF03594 | mir-3817 | MIPF0001217 |
| RF03595 | mir-3569 | MIPF0001797 |
| RF03596 | mir-5880 | MIPF0001483 |
| RF03597 | MIR6438 | MIPF0001538 |
| RF03598 | MIR6024 | MIPF0001459 |
| RF03599 | MIR1883 | MIPF0000683 |
| RF03600 | MIR3627 | MIPF0001471 |
| RF03601 | MIR7993 | MIPF0001663 |
| RF03602 | mir-8250 | MIPF0002067 |
| RF03603 | mir-4899 | MIPF0001837 |
| RF03604 | mir-5728 | MIPF0001558 |
| RF03605 | MIR4221 | MIPF0001128 |
| RF03606 | mir-2071 | MIPF0001681 |
| RF03607 | mir-9447 | MIPF0001994 |
| RF03608 | mir-9440 | MIPF0001915 |
| RF03609 | mir-5470 | MIPF0001967 |
| RF03610 | mir-H7 | MIPF0000929 |
| RF03611 | mir-7147 | MIPF0001705 |
| RF03612 | mir-608 | MIPF0000972 |
| RF03613 | MIR6234 | MIPF0001642 |
| RF03614 | mir-3431 | MIPF0002046 |
| RF03615 | mir-4877 | MIPF0001814 |
| RF03616 | MIR1888 | MIPF0001333 |
| RF03617 | MIR8757 | MIPF0002074 |
| RF03618 | mir-5592 | MIPF0001366 |
| RF03619 | mir-5729 | MIPF0001473 |
| RF03620 | mir-1233 | MIPF0000578 |
| RF03621 | MIR4245 | MIPF0001495 |
| RF03622 | mir-731 | MIPF0001354 |
| RF03623 | mir-1252 | MIPF0000968 |
| RF03624 | mir-5345 | MIPF0001357 |
| RF03625 | MIR6152 | MIPF0001607 |
| RF03626 | mir-9791 | MIPF0002118 |
| RF03627 | mir-9168 | MIPF0002095 |
| RF03628 | mir-2355 | MIPF0000938 |
| RF03629 | mir-641 | MIPF0000679 |
| RF03630 | mir-2818 | MIPF0001003 |
| RF03631 | MIR3440 | MIPF0001127 |
| RF03632 | MIR1882 | MIPF0000571 |
| RF03633 | mir-5882 | MIPF0001418 |
| RF03634 | mir-3524 | MIPF0001215 |
| RF03635 | mir-2066 | MIPF0001819 |
| RF03636 | MIR4376 | MIPF0001809 |
| RF03637 | mir-5879 | MIPF0001472 |
| RF03638 | mir-1537 | MIPF0000917 |
| RF03639 | mir-619 | MIPF0000557 |
| RF03640 | mir-3747 | MIPF0001223 |
| RF03641 | mir-991 | MIPF0000957 |
| RF03642 | MIR5743 | MIPF0001572 |
| RF03643 | mir-2056 | MIPF0001734 |
| RF03644 | mir-2064 | MIPF0001727 |
| RF03645 | mir-9080 | MIPF0001982 |
| RF03646 | MIR9776 | MIPF0001990 |
| RF03647 | mir-7640 | MIPF0001783 |
| RF03648 | mir-9413 | MIPF0002079 |
| RF03649 | MIR5643 | MIPF0001348 |
| RF03650 | mir-1290 | MIPF0000610 |
| RF03651 | mir-2070 | MIPF0001796 |
| RF03652 | mir-2329 | MIPF0000785 |
| RF03653 | mir-971 | MIPF0000995 |
| RF03654 | MIR4239 | MIPF0001164 |
| RF03655 | MIR1887 | MIPF0001125 |
| RF03656 | mir-2432 | MIPF0002055 |
| RF03657 | MIR6145 | MIPF0001620 |
| RF03658 | mir-2300 | MIPF0000770 |
| RF03659 | MIR2871 | MIPF0000986 |
| RF03660 | mir-4018 | MIPF0001259 |
| RF03661 | mir-933 | MIPF0000505 |
| RF03662 | mir-2060 | MIPF0001654 |
| RF03663 | mir-5911 | MIPF0001407 |
| RF03664 | MIR6288 | MIPF0001696 |
| RF03665 | MIR4359 | MIPF0001181 |
| RF03666 | MIR7724 | MIPF0001802 |
| RF03667 | MIR1031 | MIPF0000416 |
| RF03668 | mir-1788 | MIPF0000792 |
| RF03669 | mir-2767 | MIPF0001446 |
| RF03670 | MIR3932 | MIPF0001159 |
| RF03671 | MIR3522 | MIPF0000936 |
| RF03672 | MIR840 | MIPF0001129 |
| RF03673 | mir-2822 | MIPF0000970 |
| RF03674 | MIR5387 | MIPF0001305 |
| RF03675 | mir-1001 | MIPF0001048 |
| RF03676 | MIR5148 | MIPF0001298 |
| RF03677 | mir-1304 | MIPF0001064 |
| RF03678 | mir-3956 | MIPF0001577 |
| RF03679 | MIR2084 | MIPF0000809 |
| RF03680 | mir-9465 | MIPF0002107 |
| RF03681 | mir-1295 | MIPF0000676 |
| RF03682 | mir-3912 | MIPF0001601 |
| RF03683 | mir-462 | MIPF0001755 |
| RF03684 | mir-1990 | MIPF0000718 |
| RF03685 | MIR9677 | MIPF0001925 |
| RF03686 | mir-9424 | MIPF0002037 |
| RF03687 | mir-2011 | MIPF0000739 |
| RF03688 | mir-3045 | MIPF0001120 |
| RF03689 | mir-5892 | MIPF0001539 |
| RF03690 | mir-9058 | MIPF0002028 |
| RF03691 | mir-2765 | MIPF0001008 |
| RF03692 | MIR827 | MIPF0000726 |
| RF03693 | MIR1919 | MIPF0000582 |
| RF03694 | mir-266 | MIPF0000281 |
| RF03695 | MIR6151 | MIPF0001374 |
| RF03696 | mir-4904 | MIPF0001786 |
| RF03697 | mir-722 | MIPF0001612 |
| RF03698 | mir-2861 | MIPF0000963 |
| RF03699 | mir-729 | MIPF0001307 |
| RF03700 | mir-1923 | MIPF0000930 |
| RF03701 | mir-4041 | MIPF0001266 |
| RF03702 | MIR851 | MIPF0001168 |
| RF03703 | mir-3622 | MIPF0001179 |
| RF03704 | MIR4380 | MIPF0001117 |
| RF03705 | mir-1278 | MIPF0000591 |
| RF03706 | mir-2253 | MIPF0000915 |
| RF03707 | mir-1892 | MIPF0000799 |
| RF03709 | MIR1878 | MIPF0001218 |
| RF03710 | mir-1451 | MIPF0001049 |
| RF03711 | mir-784 | MIPF0000999 |
| RF03712 | MIR6450 | MIPF0001528 |
| RF03713 | mir-1756 | MIPF0000639 |
| RF03714 | MIR774 | MIPF0001101 |
| RF03715 | mir-977 | MIPF0000931 |
| RF03716 | mir-7131 | MIPF0001675 |
| RF03717 | mir-3119 | MIPF0000971 |
| RF03718 | mir-2804 | MIPF0000851 |
| RF03719 | mir-3914 | MIPF0001169 |
| RF03720 | MIR4228 | MIPF0001185 |
| RF03721 | mir-3478 | MIPF0002075 |
| RF03722 | MIR4378 | MIPF0001192 |
| RF03723 | MIR7125 | MIPF0001851 |
| RF03724 | mir-1013 | MIPF0000985 |
| RF03725 | mir-7094 | MIPF0001704 |
| RF03726 | mir-m21-1 | MIPF0000453 |
| RF03727 | mir-3477 | MIPF0001227 |
| RF03728 | MIR1511 | MIPF0001743 |
| RF03729 | mir-2363 | MIPF0000806 |
| RF03730 | mir-7615 | MIPF0001729 |
| RF03731 | mir-7371 | MIPF0001652 |
| RF03732 | mir-2057 | MIPF0001745 |
| RF03733 | mir-BART14 | MIPF0000959 |
| RF03734 | mir-4773 | MIPF0001250 |
| RF03735 | mir-1502 | MIPF0001213 |
| RF03736 | mir-3130 | MIPF0000845 |
| RF03737 | mir-8315 | MIPF0002039 |
| RF03738 | mir-2320 | MIPF0001253 |
| RF03739 | mir-4529 | MIPF0001451 |
| RF03740 | mir-921 | MIPF0000982 |
| RF03741 | mir-325 | MIPF0000147 |
| RF03742 | mir-2162 | MIPF0001950 |
| RF03743 | MIR7696 | MIPF0001669 |
| RF03744 | MIR3704 | MIPF0001425 |
| RF03745 | mir-2063 | MIPF0001752 |
| RF03746 | mir-2318 | MIPF0001968 |
| RF03747 | mir-4536 | MIPF0001319 |
| RF03748 | mir-2772 | MIPF0000906 |
| RF03749 | MIR2617 | MIPF0000907 |
| RF03750 | mir-2489 | MIPF0000953 |
| RF03751 | mir-7445 | MIPF0001857 |
| RF03752 | MIR5292 | MIPF0001329 |
| RF03753 | mir-1986 | MIPF0000741 |
| RF03754 | mir-3116 | MIPF0001002 |
| RF03755 | mir-4005 | MIPF0001007 |
| RF03756 | mir-4891 | MIPF0001856 |
| RF03757 | mir-6536 | MIPF0001479 |
| RF03758 | mir-2756 | MIPF0001391 |
| RF03759 | MIR8703 | MIPF0001959 |
| RF03760 | mir-2041 | MIPF0001913 |
| RF03761 | mir-5897 | MIPF0001517 |
| RF03762 | MIR859 | MIPF0001191 |
| RF03763 | mir-2568 | MIPF0001004 |
| RF03764 | mir-647 | MIPF0000939 |
| RF03765 | mir-2059 | MIPF0001683 |
| RF03766 | mir-3155 | MIPF0001242 |
| RF03767 | mir-3432 | MIPF0001187 |
| RF03768 | mir-1329 | MIPF0000635 |
| RF03769 | mir-3913 | MIPF0001134 |
| RF03770 | mir-125 | MIPF0000733 |
| RF03771 | mir-341 | MIPF0000269 |
| RF03772 | mir-4867 | MIPF0001707 |
| RF03773 | mir-978 | MIPF0000926 |
| RF03774 | mir-2110 | MIPF0001399 |
| RF03775 | mir-7578 | MIPF0001810 |
| RF03776 | mir-2068 | MIPF0001700 |
| RF03777 | MIR533 | MIPF0000356 |
| RF03778 | mir-483 | MIPF0000180 |
| RF03779 | mir-1805 | MIPF0000947 |
| RF03780 | mir-3389 | MIPF0001581 |
| RF03781 | mir-4524 | MIPF0001330 |
| RF03782 | mir-3661 | MIPF0001530 |
| RF03783 | mir-4854 | MIPF0001908 |
| RF03784 | mir-2240 | MIPF0000838 |
| RF03785 | mir-974 | MIPF0002083 |
| RF03786 | mir-9449 | MIPF0001976 |
| RF03787 | mir-2010 | MIPF0002019 |
| RF03788 | MIR5304 | MIPF0001713 |
| RF03789 | MIR5269 | MIPF0001370 |
| RF03790 | MIR8008 | MIPF0001836 |
| RF03791 | mir-4526 | MIPF0001545 |
| RF03792 | MIR6462 | MIPF0001393 |
| RF03793 | MIR810 | MIPF0000497 |
| RF03794 | MIR5298 | MIPF0001286 |
| RF03795 | MIR6201 | MIPF0001960 |
| RF03796 | mir-5984 | MIPF0001468 |
| RF03797 | mir-9235 | MIPF0001906 |
| RF03798 | MIR4227 | MIPF0001115 |
| RF03799 | mir-3202 | MIPF0000846 |
| RF03800 | mir-4477 | MIPF0001274 |
| RF03801 | mir-7143 | MIPF0002024 |
| RF03802 | MIR5649 | MIPF0001369 |
| RF03803 | MIR1886 | MIPF0001647 |
| RF03804 | mir-4989 | MIPF0001239 |
| RF03805 | bpcv-mir-B1 | MIPF0001235 |
| RF03806 | mir-2007 | MIPF0002094 |
| RF03807 | mir-4890 | MIPF0001703 |
| RF03808 | mir-3957 | MIPF0001605 |
| RF03809 | mir-1203 | MIPF0000633 |
| RF03810 | mir-8932 | MIPF0002071 |
| RF03811 | mir-1896 | MIPF0001981 |
| RF03812 | mir-BART19 | MIPF0000983 |
| RF03813 | MIR8041 | MIPF0001773 |
| RF03814 | mir-2797 | MIPF0000911 |
| RF03815 | mir-3051 | MIPF0001114 |
| RF03816 | MIR6297 | MIPF0001754 |
| RF03817 | mir-3836 | MIPF0001231 |
| RF03818 | MIR7994 | MIPF0001763 |
| RF03819 | mir-Ro6-3 | MIPF0001628 |
| RF03820 | mir-5429 | MIPF0001315 |
| RF03821 | mir-2002 | MIPF0001989 |
| RF03822 | mir-3931 | MIPF0001585 |
| RF03823 | mir-1814 | MIPF0002102 |
| RF03824 | MIR5285 | MIPF0001292 |
| RF03825 | mir-2335 | MIPF0001971 |
| RF03826 | MIR825 | MIPF0001178 |
| RF03827 | mir-2229 | MIPF0001694 |
| RF03828 | MIR831 | MIPF0001132 |
| RF03829 | mir-1257 | MIPF0000998 |
| RF03830 | mir-1003 | MIPF0000923 |
| RF03831 | mir-627 | MIPF0000550 |
| RF03832 | mir-H4 | MIPF0001247 |
| RF03833 | mir-5928 | MIPF0001443 |
| RF03834 | mir-3173 | MIPF0001383 |
| RF03835 | mir-955 | MIPF0000989 |
| RF03836 | mir-3547 | MIPF0001788 |
| RF03837 | mir-3138 | MIPF0001428 |
| RF03838 | MIR8032 | MIPF0001653 |
| RF03839 | MIR2599 | MIPF0000913 |
| RF03840 | mir-3064 | MIPF0001238 |
| RF03841 | mir-8941 | MIPF0001932 |
| RF03842 | MIR2680 | MIPF0000841 |
| RF03843 | MIR2600 | MIPF0001284 |
| RF03844 | mir-3190 | MIPF0001251 |
| RF03845 | mir-3327 | MIPF0001547 |
| RF03846 | mir-4861 | MIPF0001822 |
| RF03847 | mir-1538 | MIPF0000903 |
| RF03848 | mir-1549 | MIPF0001345 |
| RF03849 | MIR1219 | MIPF0000346 |
| RF03850 | mir-961 | MIPF0001006 |
| RF03851 | mir-1002 | MIPF0000909 |
| RF03852 | mir-610 | MIPF0000661 |
| RF03853 | mir-2013 | MIPF0001777 |
| RF03854 | mir-1949 | MIPF0001139 |
| RF03855 | MIR5241 | MIPF0001290 |
| RF03856 | mir-1188 | MIPF0001188 |
| RF03857 | mir-3129 | MIPF0001458 |
| RF03858 | mir-1991 | MIPF0000732 |
| RF03859 | mir-6132 | MIPF0001438 |
| RF03860 | MIR1520 | MIPF0000581 |
| RF03861 | mir-4484 | MIPF0001440 |
| RF03862 | MIR5274 | MIPF0001311 |
| RF03863 | mir-1258 | MIPF0000684 |
| RF03864 | mir-BART9 | MIPF0000955 |
| RF03865 | mir-7637 | MIPF0001774 |
| RF03866 | mir-1889 | MIPF0000954 |
| RF03867 | mir-4081 | MIPF0001241 |
| RF03868 | mir-H2 | MIPF0001214 |
| RF03869 | mir-2834 | MIPF0001047 |
| RF03870 | mir-2040 | MIPF0000713 |
| RF03871 | mir-5681 | MIPF0001343 |
| RF03872 | MIR6426 | MIPF0001561 |
| RF03873 | mir-3192 | MIPF0001597 |
| RF03874 | mir-8834 | MIPF0002099 |
| RF03875 | MIR5272 | MIPF0001278 |
| RF03876 | mir-7132 | MIPF0002098 |
| RF03877 | mir-3937 | MIPF0001500 |
| RF03878 | mir-B6 | MIPF0001260 |
| RF03879 | mir-2831 | MIPF0000978 |
| RF03880 | mir-1684 | MIPF0001537 |
| RF03881 | mir-59 | MIPF0000940 |
| RF03882 | mir-4889 | MIPF0001686 |
| RF03883 | mir-9256 | MIPF0001895 |
| RF03884 | mir-725 | MIPF0001355 |
| RF03885 | mir-4178 | MIPF0000928 |
| RF03886 | mir-1993 | MIPF0000716 |
| RF03887 | mir-2511 | MIPF0000977 |
| RF03888 | mir-3160 | MIPF0001028 |
| RF03889 | mir-4776 | MIPF0001210 |
| RF03890 | mir-B8 | MIPF0002021 |
| RF03891 | mir-2545 | MIPF0000994 |
| RF03892 | mir-4860 | MIPF0001765 |
| RF03893 | mir-4879 | MIPF0001780 |
| RF03894 | mir-5736 | MIPF0001480 |
| RF03895 | MIR2679 | MIPF0000892 |
| RF03896 | MIR2275 | MIPF0000797 |
| RF03897 | mir-4077 | MIPF0000905 |
| RF03898 | mir-7616 | MIPF0001718 |
| RF03899 | mir-5912 | MIPF0001497 |
| RF03900 | mir-9242 | MIPF0002032 |
| RF03901 | mir-BART18 | MIPF0000948 |
| RF03902 | MIR7784 | MIPF0001728 |
| RF03903 | mir-679 | MIPF0001564 |
| RF03904 | MIR7991 | MIPF0001701 |
| RF03905 | MIR6149 | MIPF0001580 |
| RF03906 | mir-3199 | MIPF0001026 |
| RF03907 | mir-4428 | MIPF0001630 |
| RF03908 | MIR6459 | MIPF0001730 |
| RF03909 | mir-3804 | MIPF0001255 |
| RF03910 | mir-H5 | MIPF0000966 |
| RF03911 | mir-1910 | MIPF0000987 |
| RF03912 | mir-4659 | MIPF0001256 |
| RF03913 | mir-992 | MIPF0000914 |
| RF03914 | mir-BART21 | MIPF0000961 |
| RF03915 | MIR5286 | MIPF0001342 |
| RF03916 | MIR400 | MIPF0001157 |
| RF03917 | mir-4449 | MIPF0002103 |
| RF03918 | mir-4922 | MIPF0001229 |
| RF03919 | mir-1540 | MIPF0001772 |
| RF03920 | mir-237 | MIPF0000904 |
| RF03921 | MIR860 | MIPF0001176 |
| RF03922 | MIR2119 | MIPF0000762 |
| RF03923 | mir-964 | MIPF0000949 |
| RF03924 | mir-1228 | MIPF0000924 |
| RF03925 | mir-1011 | MIPF0000969 |
| RF03926 | MIR1435 | MIPF0000760 |
| RF03927 | MIR538 | MIPF0000161 |
| RF03929 | mir-6579 | MIPF0001492 |
| RF03930 | mir-1909 | MIPF0000973 |
| RF03931 | mir-2061 | MIPF0001790 |
| RF03932 | mir-1597 | MIPF0000764 |
| RF03933 | MIR6445 | MIPF0001621 |
| RF03934 | mir-m107-1 | MIPF0000446 |
| RF03935 | mir-763 | MIPF0000722 |
| RF03936 | mir-4079 | MIPF0001273 |
| RF03937 | mir-9456 | MIPF0001974 |
| RF03938 | mir-4662 | MIPF0001245 |
| RF03939 | mir-4868 | MIPF0001233 |
| RF03940 | MIR9563 | MIPF0001941 |
| RF03941 | mir-3158 | MIPF0001065 |
| RF03942 | mir-5727 | MIPF0001627 |
| RF03943 | mir-1824 | MIPF0001717 |
| RF03944 | mir-935 | MIPF0000606 |
| RF03945 | mir-4876 | MIPF0001776 |
| RF03946 | mir-3132 | MIPF0001509 |
| RF03947 | mir-658 | MIPF0000643 |
| RF03948 | mir-5045 | MIPF0001341 |
| RF03949 | mir-8830 | MIPF0001924 |
| RF03950 | mir-4928 | MIPF0001818 |
| RF03951 | MIR7725 | MIPF0001792 |
| RF03952 | mir-3718 | MIPF0001225 |
| RF03953 | mir-5350 | MIPF0001367 |
| RF03954 | mir-8993 | MIPF0002018 |
| RF03955 | MIR2934 | MIPF0001269 |
| RF03956 | mir-737 | MIPF0001689 |
| RF03957 | mir-5856 | MIPF0001503 |
| RF03958 | mir-959 | MIPF0000910 |
| RF03959 | mir-303 | MIPF0001013 |
| RF03960 | mir-6644 | MIPF0001504 |
| RF03961 | mir-9033 | MIPF0002087 |
| RF03962 | mir-958 | MIPF0000912 |
| RF03963 | mir-2996 | MIPF0001023 |
| RF03964 | mir-H1 | MIPF0000738 |
| RF03965 | mir-8927 | MIPF0001979 |
| RF03966 | mir-76 | MIPF0000902 |
| RF03967 | mir-1270 | MIPF0000877 |
| RF03968 | mir-8934 | MIPF0002010 |
| RF03969 | mir-622 | MIPF0000619 |
| RF03970 | mir-2766 | MIPF0001608 |
| RF03971 | mir-3572 | MIPF0001244 |
| RF03972 | mir-3150 | MIPF0001102 |
| RF03973 | mir-5615 | MIPF0001318 |
| RF03974 | MIR838 | MIPF0001184 |
| RF03975 | mir-5881 | MIPF0001420 |
| RF03976 | MIR898 | MIPF0000378 |
| RF03977 | mir-976 | MIPF0000916 |
| RF03978 | mir-1269 | MIPF0000984 |
| RF03979 | mir-2006 | MIPF0001936 |
| RF03980 | mir-2012 | MIPF0000720 |
| RF03981 | mir-8799 | MIPF0001881 |
| RF03982 | mir-4865 | MIPF0001750 |
| RF03983 | MIR5284 | MIPF0001277 |
| RF03984 | MIR2933 | MIPF0001072 |
| RF03985 | mir-9250 | MIPF0002073 |
| RF03986 | mir-6095 | MIPF0001433 |
| RF03987 | MIR6136 | MIPF0001741 |
| RF03988 | mir-6301 | MIPF0001556 |
| RF03989 | mir-7950 | MIPF0001678 |
| RF03990 | mir-4888 | MIPF0001724 |
| RF03991 | mir-9209 | MIPF0001927 |
| RF03992 | mir-6302 | MIPF0001398 |
| RF03993 | mir-8536 | MIPF0002030 |
| RF03994 | mir-7558 | MIPF0001716 |
| RF03995 | mir-7957 | MIPF0001668 |
| RF03996 | mir-3012 | MIPF0001018 |
| RF03997 | mir-8510 | MIPF0001886 |
| RF03998 | mir-2160 | MIPF0000769 |
| RF03999 | mir-7883 | MIPF0001849 |
| RF04000 | MIR8139 | MIPF0001665 |
| RF04001 | mir-7918 | MIPF0001655 |
| RF04002 | MIR6143 | MIPF0001685 |
| RF04003 | mir-4841 | MIPF0001264 |
| RF04004 | mir-7940 | MIPF0001719 |
| RF04005 | mir-4599 | MIPF0001252 |
| RF04006 | mir-9338 | MIPF0001921 |
| RF04007 | mir-8489 | MIPF0002034 |
| RF04008 | mir-4880 | MIPF0001833 |
| RF04009 | mir-754 | MIPF0000749 |
| RF04010 | mir-3003 | MIPF0000818 |
| RF04011 | mir-8515 | MIPF0002064 |
| RF04012 | MIR7532 | MIPF0001820 |
| RF04013 | mir-4818 | MIPF0001211 |
| RF04014 | mir-8338 | MIPF0002060 |
| RF04015 | MIR946 | MIPF0000427 |
| RF04016 | mir-4873 | MIPF0001806 |
| RF04017 | mir-4864 | MIPF0001702 |
| RF04018 | mir-5316 | MIPF0001364 |
| RF04019 | MIR950 | MIPF0000407 |
| RF04020 | mir-4859 | MIPF0001784 |
| RF04021 | mir-3492 | MIPF0001622 |
| RF04022 | mir-4863 | MIPF0001778 |
| RF04023 | MIR1314 | MIPF0001609 |
| RF04024 | mir-9571 | MIPF0002068 |
| RF04025 | mir-7964 | MIPF0001749 |
| RF04026 | mir-8499 | MIPF0002029 |
| RF04027 | mir-1422 | MIPF0000565 |
| RF04028 | mir-7948 | MIPF0001695 |
| RF04029 | mir-8521 | MIPF0001871 |
| RF04030 | mir-6096 | MIPF0001436 |
| RF04031 | mir-2686 | MIPF0000927 |
| RF04032 | MIR2912 | MIPF0000778 |
| RF04033 | mir-8512 | MIPF0002076 |
| RF04035 | mir-7910 | MIPF0001840 |
| RF04036 | mir-2076 | MIPF0001795 |
| RF04037 | mir-7911 | MIPF0001858 |
| RF04038 | mir-5985 | MIPF0001491 |
| RF04039 | mir-2242 | MIPF0000945 |
| RF04040 | mir-4866 | MIPF0001756 |
| RF04041 | MIR6154 | MIPF0001560 |
| RF04042 | mir-7961 | MIPF0001832 |
| RF04043 | mir-2062 | MIPF0001753 |
| RF04044 | mir-7939 | MIPF0001646 |
| RF04045 | mir-2147 | MIPF0000751 |
| RF04046 | mir-2154 | MIPF0000781 |
| RF04047 | mir-9228 | MIPF0001876 |
| RF04048 | mir-7927 | MIPF0001660 |
| RF04049 | MIR7127 | MIPF0001408 |
| RF04050 | mir-2072 | MIPF0001697 |
| RF04051 | mir-8517 | MIPF0002077 |
| RF04052 | mir-466 | MIPF0000208 |
| RF04053 | mir-95 | MIPF0000098 |
| RF04054 | mir-2944 | MIPF0001335 |
| RF04055 | mir-1261 | MIPF0000802 |
| RF04056 | mir-982 | MIPF0000843 |
| RF04057 | MIR1432 | MIPF0001063 |
| RF04059 | MIR5020 | MIPF0001226 |
| RF04060 | mir-703 | MIPF0000761 |
| RF04061 | mir-1420 | MIPF0000574 |
| RF04062 | MIR949 | MIPF0001455 |
| RF04063 | MIR2629 | MIPF0000824 |
| RF04064 | MIR5267 | MIPF0001276 |
| RF04065 | mir-7563 | MIPF0001709 |
| RF04066 | mir-1256 | MIPF0000594 |
| RF04067 | mir-4520 | MIPF0001272 |
| RF04068 | mir-3927 | MIPF0001462 |
| RF04069 | MIR6135 | MIPF0001644 |
| RF04070 | MIR6440 | MIPF0001406 |
| RF04071 | MIR2863 | MIPF0000925 |
| RF04072 | mir-7386 | MIPF0001658 |
| RF04073 | mir-727 | MIPF0001371 |
| RF04074 | mir-1199 | MIPF0001659 |
| RF04075 | mir-1545 | MIPF0001711 |
| RF04076 | mir-H11 | MIPF0000833 |
| RF04077 | mir-1245 | MIPF0000620 |
| RF04078 | mir-1905 | MIPF0000753 |
| RF04079 | MIR161 | MIPF0000455 |
| RF04080 | mir-762 | MIPF0000624 |
| RF04081 | mir-1992 | MIPF0000705 |
| RF04082 | mir-54 | MIPF0000874 |
| RF04083 | MIR528 | MIPF0000868 |
| RF04084 | mir-595 | MIPF0000651 |
| RF04085 | mir-1271 | MIPF0000483 |
| RF04087 | MIR5565 | MIPF0001279 |
| RF04088 | MIR812 | MIPF0000345 |
| RF04089 | mir-753 | MIPF0000752 |
| RF04090 | mir-6529 | MIPF0001771 |
| RF04091 | mir-72 | MIPF0000277 |
| RF04092 | MIR1508 | MIPF0000719 |
| RF04093 | mir-1784 | MIPF0001873 |
| RF04094 | MIR537 | MIPF0000207 |
| RF04095 | MIR9746 | MIPF0001884 |
| RF04096 | mir-4433 | MIPF0001826 |
| RF04097 | MIR2630 | MIPF0000814 |
| RF04098 | MIR9471 | MIPF0002045 |
| RF04099 | MIR837 | MIPF0001155 |
| RF04100 | mir-493 | MIPF0000230 |
| RF04101 | mir-8864 | MIPF0001919 |
| RF04102 | MIR6457 | MIPF0001534 |
| RF04103 | mir-8356 | MIPF0001997 |
| RF04104 | MIR918 | MIPF0000374 |
| RF04105 | mir-3165 | MIPF0001635 |
| RF04106 | mir-1303 | MIPF0000608 |
| RF04107 | mir-5391 | MIPF0001944 |
| RF04108 | mir-2513 | MIPF0000980 |
| RF04109 | mir-1305 | MIPF0000965 |
| RF04110 | MIR5084 | MIPF0002026 |
| RF04111 | MIR6274 | MIPF0001706 |
| RF04112 | MIR5512 | MIPF0001478 |
| RF04113 | MIR1523 | MIPF0001327 |
| RF04114 | MIR4374 | MIPF0001152 |
| RF04115 | mir-6130 | MIPF0001413 |
| RF04116 | MIR9672 | MIPF0001942 |
| RF04117 | MIR5200 | MIPF0001904 |
| RF04118 | MIR9481 | MIPF0002096 |
| RF04119 | MIR1858 | MIPF0000650 |
| RF04120 | MIR5998 | MIPF0001613 |
| RF04121 | mir-1289 | MIPF0000626 |
| RF04122 | mir-1204 | MIPF0000671 |
| RF04123 | mir-3127 | MIPF0001439 |
| RF04124 | MIR8565 | MIPF0001889 |
| RF04125 | MIR9408 | MIPF0002027 |
| RF04126 | mir-9198 | MIPF0001887 |
| RF04127 | MIR8007 | MIPF0001839 |
| RF04128 | MIR8562 | MIPF0001861 |
| RF04129 | MIR4240 | MIPF0001198 |
| RF04130 | MIR7996 | MIPF0001691 |
| RF04131 | mir-2032 | MIPF0000736 |
| RF04132 | mir-4436 | MIPF0001236 |
| RF04133 | MIR2646 | MIPF0001010 |
| RF04134 | MIR7504 | MIPF0001798 |
| RF04135 | MIR5062 | MIPF0001922 |
| RF04136 | MIR1862 | MIPF0000580 |
| RF04137 | MIR834 | MIPF0001183 |
| RF04138 | mir-9243 | MIPF0001926 |
| RF04139 | MIR841 | MIPF0001112 |
| RF04140 | mir-3118 | MIPF0001928 |
| RF04141 | MIR8622 | MIPF0001931 |
| RF04142 | mir-1299 | MIPF0000625 |
| RF04143 | MIR869 | MIPF0001167 |
| RF04144 | MIR9783 | MIPF0001947 |
| RF04145 | MIR8706 | MIPF0001953 |
| RF04146 | mir-4742 | MIPF0001552 |
| RF04147 | MIR6224 | MIPF0001435 |
| RF04148 | MIR844 | MIPF0001197 |
| RF04149 | MIR447 | MIPF0000170 |
| RF04150 | MIR2655 | MIPF0000813 |
| RF04151 | MIR4387 | MIPF0001109 |
| RF04152 | mir-3618 | MIPF0001710 |
| RF04153 | MIR856 | MIPF0001162 |
| RF04154 | mir-1277 | MIPF0001937 |
| RF04155 | mir-9236 | MIPF0001905 |
| RF04156 | MIR2670 | MIPF0000855 |
| RF04157 | MIR9486 | MIPF0002054 |
| RF04158 | mir-568 | MIPF0000408 |
| RF04159 | MIR1437 | MIPF0001762 |
| RF04160 | MIR868 | MIPF0001173 |
| RF04161 | mir-3688 | MIPF0001263 |
| RF04162 | MIR9662 | MIPF0002119 |
| RF04163 | mir-8364 | MIPF0001877 |
| RF04164 | MIR1507 | MIPF0000699 |
| RF04165 | mir-2483 | MIPF0001532 |
| RF04166 | mir-2278 | MIPF0001522 |
| RF04167 | mir-1972 | MIPF0001025 |
| RF04168 | MIR839 | MIPF0001143 |
| RF04169 | mir-2235 | MIPF0000825 |
| RF04170 | MIR7723 | MIPF0001739 |
| RF04171 | MIR1509 | MIPF0000771 |
| RF04172 | MIR8675 | MIPF0002106 |
| RF04173 | MIR847 | MIPF0001171 |
| RF04174 | MIR4208 | MIPF0001108 |
| RF04175 | mir-1260b | MIPF0001381 |
| RF04176 | mir-3149 | MIPF0001935 |
| RF04185 | MIR2118_2 | MIPF0001409 |
| RF04186 | mir-278_2 | MIPF0000728 |
| RF04187 | MIR862 | MIPF0001145 |
| RF04193 | mir-51 | MIPF0000268 |
| RF04194 | mir-57 | MIPF0000271 |
| RF04195 | MIR6217 | MIPF0001550 |
| RF04196 | MIR5638 | MIPF0001350 |
| RF04197 | mir-9457 | MIPF0002122 |
| RF04198 | MIR8670 | MIPF0002041 |
| RF04199 | MIR6027 | MIPF0001424 |
| RF04200 | mir-9437 | MIPF0001943 |
| RF04201 | MIR7510 | MIPF0001684 |
| RF04202 | MIR5380 | MIPF0001331 |
| RF04203 | MIR1144 | MIPF0000432 |
| RF04204 | mir-3473 | MIPF0001230 |
| RF04205 | MIR2606 | MIPF0000888 |
| RF04206 | MIR9555 | MIPF0002081 |
| RF04207 | MIR5048 | MIPF0001453 |
| RF04208 | MIR8742 | MIPF0002016 |
| RF04209 | MIR7533 | MIPF0001834 |
| RF04210 | MIR8643 | MIPF0002008 |
| RF04211 | MIR5185 | MIPF0001246 |
| RF04212 | mir-8904 | MIPF0001977 |
| RF04213 | MIR7807 | MIPF0002091 |
| RF04214 | MIR7526 | MIPF0001651 |
| RF04215 | mir-9214 | MIPF0002108 |
| RF04216 | mir-509 | MIPF0000130 |
| RF04217 | mir-297 | MIPF0000204 |
| RF04223 | MIR2619 | MIPF0001294 |
| RF04224 | mir-9229 | MIPF0001875 |
| RF04225 | MIR7502 | MIPF0001869 |
| RF04226 | mir-9186 | MIPF0001868 |
| RF04227 | mir-9215 | MIPF0001893 |
| RF04228 | MIR6140 | MIPF0001666 |
| RF04229 | mir-9279 | MIPF0001911 |
| RF04230 | mir-9261 | MIPF0001938 |
| RF04231 | mir-9191 | MIPF0001987 |
| RF04232 | mir-9318 | MIPF0002056 |
| RF04233 | mir-680 | MIPF0000338 |
| RF04234 | mir-1421 | MIPF0000564 |
| RF04235 | mir-242_2 | MIPF0000731 |
| RF04236 | mir-1285 | MIPF0000559 |
| RF04237 | mir-1490 | MIPF0000525 |
| RF04238 | mir-4012 | MIPF0000964 |
| RF04239 | mir-4679 | MIPF0001228 |
| RF04240 | mir-4716 | MIPF0001476 |
| RF04241 | mir-5014 | MIPF0001634 |
| RF04242 | mir-5595 | MIPF0001590 |
| RF04243 | MIR169_3 | MIPF0000253 |
| RF04244 | MIR169_6 | MIPF0001058 |
| RF04245 | MIR169_7 | MIPF0001690 |
| RF04246 | MIR169_8 | MIPF0001712 |
| RF04248 | MIR7486 | MIPF0001682 |
| RF04249 | mir-310 | MIPF0000566 |
| RF04250 | mir-236 | MIPF0000232 |
| RF04251 | MIR5070 | MIPF0002013 |
| RF04252 | mir-8186 | MIPF0002035 |
| RF04254 | mir-1677 | MIPF0000849 |
| RF04255 | MIR8001 | MIPF0001852 |
| RF04257 | mir-9230 | MIPF0002035 |
| RF04258 | mir-994 | MIPF0001045 |
| RF04259 | mir-2003 | MIPF0002049 |
| RF04260 | mir-2005 | MIPF0001956 |
| RF04261 | MIR1861 | MIPF0000567 |
| RF04262 | MIR1319 | MIPF0001448 |
| RF04263 | MIR2593 | MIPF0000733 |
| RF04264 | mir-5408 | MIPF0001313 |
| RF04265 | mir-4650 | MIPF00001234 |
| RF04266 | mir-9412 | MIPF0001891 |
| RF04267 | mir-362 | MIPF0000209 |
| RF04268 | mir-512 | MIPF0000518 |
| RF04269 | mir-373 | MIPF0000500 |
| RF04270 | mir-743 | MIPF0000386 |
| RF04271 | mir-1244 | MIPF0000569 |
| RF04272 | mir-8908 | MIPF0001890 |
| RF04273 | MIR827_2 | MIPF0001769 |
| RF04274 | mir-9223 | MIPF0001885 |
| RF04275 | mir-2733 | MIPF0000766 |
| RF04276 | MIR4371 | MIPF0001141 |
| RF04277 | mir-1268 | MIPF0000946 |
| RF04278 | MIR4372 | MIPF0001257 |
| RF04279 | mir-H20 | MIPF0001760 |
| RF04280 | mir-890 | MIPF0000386 |
| RF04281 | mir-8791 | MIPF0001882 |
| RF04282 | mir-507 | MIPF0000130 |
| RF04283 | mir-3135 | MIPF0001219 |
| RF04284 | mir-532 | MIPF0000113 |
| RF04285 | mir-660 | MIPF0000113 |
| RF04286 | mir-513 | MIPF0000130 |
| RF04287 | mir-9195 | MIPF0001939 |
| RF04288 | mir-9201 | MIPF0001898 |
| RF04289 | mir-3596 | MIPF0001194 |
| RF04290 | mir-1197 | MIPF0000126 |
| RF04291 | mir-368 | MIPF0000091 |
| RF04292 | mir-379 | MIPF0000126 |
| RF04293 | mir-889 | MIPF0000514 |
| RF04294 | mir-3578 | MIPF0001166 |
| RF04295 | mir-329 | MIPF0000110 |
| RF04296 | mir-485 | MIPF0000201 |
| RF04297 | mir-35_2 | MIPF0001648 |
| RF04298 | mir-36_2 | MIPF0001664 |
| RF04299 | MIR814 | MIPF0000351 |
| RF04300 | mir-39 | MIPF0000304 |
| RF04301 | mir-200 | MIPF0000491 |
| RF04302 | mir-506 | MIPF0000130 |
| RF04303 | MIR162_2 | MIPF0000169 |

